# Identification and antimicrobial properties of bacteria isolated from naturally decaying wood

**DOI:** 10.1101/2020.01.07.896464

**Authors:** Tanja R. Scheublin, Anna M. Kielak, Marlies van den Berg, Johannes A. van Veen, Wietse de Boer

## Abstract

Research on wood decay in forest ecosystems has traditionally focused on wood-rot fungi, which lead the decay process through attack of the lignocellulose complex. The role of bacteria, which can be highly abundant, is still unclear. Wood-inhabiting bacteria are thought to be nutritionally dependent on decay activities of wood-rot fungi. Therefore, we hypothesized that these bacteria are not antagonistic against wood-rot fungi whereas antagonistic activity against other bacteria may be high (resource competition). This was examined for decaying wood in temperate forests. We found that the abundance of cultivable bacteria in decaying wood can be highly variable. The general pattern is an increase of bacteria with progressive decay, but we also identified several fungi that were apparently able to exclude bacteria from their woody territory. We established a bacterial collection which is highly representative for decaying wood with typical wood-inhabiting taxa: *Xanthomonadaceae*, *Acetobacteraceae*, *Caulobacteraceae*, *Methylovirgula*, *Sphingomonas*, *Burkholderia* and *Granulicella*. *In vitro* antagonistic activity against other bacteria and fungi was evaluated. In contrast to our hypothesis, we found surprisingly low antagonistic activity against bacteria (<2% of isolates), while antagonism against fungi was more prevalent. These results may point at a prominent role of mycophagy (growth at the expense of living fungi) among wood-inhabiting bacteria.

## Introduction

The process of wood decay has been extensively studied; both from the perspective of functioning of forest ecosystems (e.g. carbon and nutrient cycling) and of wood industry and construction (Schmidt 2006; van der Wal *et al* 2013). The main players in aerobically decaying wood are wood-rot fungi. They lead the process of wood decay with their ability to attack the lignocellulose complex using a complex machinery of extracellular enzymes and chemical mediators (Dashtban *et al* 2010). The presence of bacteria in wood decayed by wood-rot fungi has also been demonstrated, though information on abundance, identity and functional characteristics is still very limited (Johnston *et al* 2016). The contribution of bacteria to wood decay via direct attack of the ligno-cellulose complex is assumed to be limited to circumstances where fungal decay is suppressed, like in water-logged wood and foundation piles (Klaassen 2008). However, the indirect influence of bacteria on wood decay via mutualistic or competitive interactions with wood-rot fungi is potentially large (de Boer and van der Wal 2008; Johnston *et al* 2016).

Information on the bacterial community composition in decaying wood is increasingly becoming available (Johnston et al 2016). Bacterial taxa that are commonly found in decaying wood include *Acidobacteria*, Alpha-, Beta- and Gamma*proteobacteria* (*Beijerinckiaceae*, *Burkholderiaceae*, *Xanthomonadaceae*) and *Actinobacteria* (Hervé *et al* 2014; Hoppe *et al* 2015; Kielak *et al* 2016; Rina-Kanto *et al* 2016; Tláskal *et al* 2017; Probst *et al* 2018). Bacterial biomass and diversity increases with progressive decay (Hoppe et al 2015; Kielak et al 2016; Tláskal *et al* 2017; Probst *et al* 2018) and the relative abundance of some bacterial taxa is associated with the stage of decay, e.g. the genus *Methylovirgula* is generally found in intermediate to advanced stages. Besides decay stage, tree species is also influencing the bacterial community composition (Hoppe et al 2015). No relationship between bacterial and fungal communities was found in a pine decay range (Kielak et al 2016), indicating that the physico-chemical parameters of the wood rather than the composition of fungi were the main determinants for bacterial community composition. In contrast, co-occurrence patterns between fungi and nitrogen fixing bacteria were found in decaying *Fagus* and *Picea* wood (Hoppe et al 2014).

It is well known that bacteria and fungi can have a large impact on each other’s growth and performance via the production of secondary metabolites, but these interactions have hardly been investigated in the context of wood decay. In contrast, antagonistic interactions among different wood decaying fungi have received a lot of attention (Boddy 2000; Hiscox *et al* 2018). Wood decaying fungi heavily compete for substrate and a wide range of antimicrobial compounds have been identified. These compounds can inhibit fungi as well as bacteria (Suay et al 2000; Barros et al 2008). In addition, acidification of wood by fungal production of organic acids, like oxalic acid, have a big impact on the growth conditions for bacteria. Indeed, a sharp decline in the number of bacteria in woodblocks on forest soil has been observed after invasion of the blocks by the white-rot fungus *Hypholoma fasciculare* and rapid acidification rather than toxic compounds were the most likely cause of this decline (Folman et al 2008; de Boer et al 2010). In contrast, high bacterial numbers of up to 10^10^ colony forming units per gram of wood have been observed in naturally acidic, decaying wood covered with *H. fasciculare* fruiting bodies (Valaskova et al 2009). Currently, it is still unclear, how bacterial abundance and functioning is influenced by wood-rot fungi during the decay process.

Conversely, bacteria in decaying wood can also influence wood-rot fungi. Co-inoculation studies with wood decaying fungi and bacteria have reported positive as well as negative effects on the decay process (Blanchette and Shaw 1978; Murray and Woodard 2003). Only one study examined the direct interactions between wood-rot fungi and co-existing bacteria and reported neutral, inhibitory and in one case stimulatory effects on fungal growth (Kamei et al 2012). No studies exist on antibacterial properties of bacteria from decaying wood. We hypothesize that their combative strategies are primarily directed towards other bacteria, while leaving the wood-rot fungi unaffected. This hypothesis is based on the assumption that bacteria in non-saturated decaying wood are nutritionally dependent on the release of oligomers of wood polymers by decay fungi (de Boer and van der Wal 2008). To test this hypothesis we aimed to get a collection of bacterial strains that is representative for different stages of decaying wood in a temperate forest. Of the identified bacterial strains, antibacterial and antifungal properties were determined by *in vitro* screenings. In addition to identification of antimicrobial properties, we also determined abundance of cultivable bacteria in wood samples with different wood-rot fungi and in different decay stages.

## Materials and methods

### Sample collection

Decaying wood samples were collected in a mixed forest close to the village of Wolfheze in the centre of The Netherlands (51°59’39’’N; 5°47’39’’E). For a pilot experiment, wood samples from decaying birch trees with either *Piptoporus betulinus*, *Fomes fomentarius*, *Stereum subtomentosum* or *Plicaturopsis crispa* fruiting bodies were collected in duplo in autumn 2012. In addition, two coniferous samples (Abies fir and pine wood) with clear visual indications of brown-rot patterns were collected in the same year. For the main experiment, twenty samples of pine wood (*Pinus sylvestris*) in different stages of decay were collected in autumn 2013. For all samples, the bark was removed and slices of wood were surface sterilized under UV light for 30 min. Saw dust was produced with a wood drill, sterilized with 70% ethanol. Highly decayed wood samples, which could not be drilled, were fragmented using sterile forceps and scalpel. Frozen saw dust samples from oak stumps two and five years after cutting (van der Wal *et al* 2015) were included in the pilot.

### Bacterial enumeration and identification

About 0.4 g of fresh saw dust was weighed into a 5 mL tube and mixed with 4 mL 0.9% sodium chloride buffered at pH 5.0 with 1.95 g 2-(*N*-morpholino)ethanesulfonic acid (MES) per liter. Tubes were shaken at 300 rpm for 90 min, sonicated twice for 30 seconds and shaken for another 30 min. For the pilot experiments, two times 50 μL of 10^0^, 10^−1^, 10^−2^, 10^−3^, 10^−4^ and 10^−5^ dilutions were spread on water-yeast agar pH 5.0 with cycloheximide (containing 1 g sodium chloride, 0.1 g yeast extract, 1.95 g MES, 20 g agar and 100 mg cycloheximide per liter). Previous experiments identified this low-nutrient, low-pH medium as the most suitable medium for the isolation of bacteria from decaying wood (Valášková *et al* 2009). Plates were incubated at 20 °C and bacterial colonies were counted after 5 weeks of growth. A maximum of 24 colonies per wood sample were randomly collected for identification by 16S rDNA sequencing. For the main experiment, two times 50 μL of 10^−2^, 10^−3^, 10^−4^, 10^−5^ and 10^−6^ dilutions were spread on water-yeast agar pH 5.0 with cycloheximide, thiabendazole and methanol (containing 1 g sodium chloride, 0.1 g yeast extract, 1.95 g MES, 20 g agar, 100 mg cycloheximide, 11 mg thiabendazole and 1 g methanol per liter). Methanol was included, because in the pilot experiments, diminished growth was observed for *Methylovirgula* isolates in the absence of methanol. The applied concentration of methanol is unlikely to be lethal for the majority of bacteria (Wadhwani *et al* 2008). A combination of cycloheximide and thiabendazole appeared to be more effective to inhibit fungal growth that cycloheximide alone (Hol *et al* 2015). Plates were incubated at 20 °C and bacterial colonies were counted after 10 days, 3 weeks and 6 weeks. At each time point, a maximum of 24 colonies per wood sample were collected for further identification.

Bacteria were identified by 16S rDNA sequencing. Colony PCR was performed with primers 27f (5’-GAGTTTGATCMTGGCTCAG-3’) and 1492r (5’-GRTACCTTGTTACGACTT-3’; Lane, 1991). The PCR mixture contained 0.04 U FastStart Taq DNA polymerase, 1× buffer (Roche Diagnostics, Mannheim, Germany), 0.6 μM of each primer and 200 μM of each deoxynucleoside triphosphate in a total volume of 25 μL. The PCR cycling regime was (1) one cycle of 5 min at 94°C, (2) 35 cycles of 1 min at 94°C, 1 min at 55°C, and 1.5 min at 72°C, and (3) one final extension cycle of 10 min at 72°C. PCR products were verified by agarose gel electrophoresis. Fragments were sequenced (Macrogen, Seoul, Korea) from both directions in the pilot experiment and with primer 1492r in the main experiment.

Low quality regions of 16S rDNA sequences were trimmed and DNA fragments that were sequenced from both sides were assembled (DNA Baser Sequence Assembler; www.dnabaser.com). Sequences were identified using the EzTaxon server (http://www.ezbiocloud.net/eztaxon; Kim et al 2012) and classified into a taxonomic group with RDP Classifier (version 2.10, training set 14; Wang et al 2007). Trimmed and assembled sequences were deposited in GenBank under accession numbers KY907704-KY908306. The relative abundance of taxa was calculated, including only randomly picked colonies and taking into account the relative abundance at each time point as derived from colony counts. Bacterial communities from the same wood samples have been previously identified with pyrosequencing (Kielak *et al* 2016). Abundance of bacterial taxa as derived from sequenced isolates was plotted against the relative abundance in the pyrosequencing data for comparison. This was done at class as well as genus level.

For a comparative analysis, sequences from two previously published studies were recovered from GenBank (Folman *et al* 2008, Valášková *et al* 2009) and reclassified with RDP Classifier version 2.10 and training set 14. Presence of bacterial taxa was compared between the two published studies and the present work.

### Physicochemical characteristics

Wood density was determined by water displacement of wood blocks (Olesen 1971). Moisture content was determined by drying wood dust at 60 °C for 4 days. Dried wood was grinded with liquid nitrogen and carbon and nitrogen content were measured on a Flash EA1112 CN analyzer (Interscience, Breda, the Netherlands).

Water extracts of saw dust were prepared by shaking 0.3 g of fresh saw dust with 6 mL milli-Q water at 300 rpm for 1 hour. Water extracts were used to determine pH, manganese peroxidase, laccase and cellulase activities as previously described (Valášková *et al* 2009). Ergosterol content was determined as an indication for fungal biomass with alkaline extraction and HPLC analysis (Baath 2001; de Ridder-Duine *et al* 2006).

A correlation matrix was produced for all wood physicochemical parameters (density, moisture content, pH, C/N ratio), fungal biomass (ergosterol content), enzyme activities (Mn peroxidase, laccase, cellulase) and bacterial numbers (CFU per gram dry wood after 6 weeks). Correlations were calculated with Kendall’s tau-b and Spearmann’s rho coefficients using SPSS version 23.0.

### Antibacterial activity screening

A total of 635 isolates from decaying pine wood (main experiment) were screened for antibacterial activity. They were routinely grown in 24 well plates on water-yeast agar pH5 with 1 g L^−1^ methanol (WYA5m) at 20 °C. The bacteria were transferred to 1-well plates (Greiner Bio-One; www.gbo.com) containing 35 mL WYA5m with the use of a custom-made 24-pin tool. The plates were incubated at 20 °C for 1, 2 and 5 weeks for the isolates collected after 10 days, 3 weeks and 6 weeks respectively. Antibacterial activity was tested against four bacterial strains, *Escherichia coli* WA123, *Staphylococcus aureus* 533R4, *Dyella* strain WH32 and *Burkholderia* strain 4-A6. *E. coli* and *S. aureus* are clinically relevant strains for the development of new antibiotics and the *Dyella* and *Burkholderia* strains are ecologically relevant, as they originate from decaying wood. *E. coli* and *S. aureus* were routinely grown at 37 °C in Luria broth (LB) and *Dyella* and *Burkholderia* at 25 °C in 1/10 tryptic soy broth pH5 (TSA5, containing 3 g tryptic soy broth (Oxoid) and 1.95 g MES per liter). Bacteria were grown overnight and the optical density at 600 nm (OD_600_) was determined. Agar medium (LB or TSA5 with 15 gram agar per liter) at 50 °C was inoculated with the bacteria to a final OD_600_ of 0.002, 0.002, 0.004 and 0.008 for *E. coli*, *S. aureus*, *Dyella* and *Burkholderia* respectively. Plates were overlaid with 15 mL and the occurrence of clear zones around the bacterial isolates was observed after one day of growth. Isolates that showed inhibition zones were streaked for purity and the overlay assay was repeated in order to verify the results of the screening.

### Antimicrobial activity of selected isolates

Thirty-six isolates were selected for a more in-depth characterization of antimicrobial properties (Suppl. Table 1). The selection was based on two criteria, (1) the isolates originated from pine samples colonized by *Stereum sanguinolentum* or *Mycena galericulata* (Kielak *et al* 2016), and (2) the isolates belonged to bacterial groups that are commonly isolated from decaying wood. Fungal strains *S. sanguinolentum* CBS 927.72 and *M. galericulata* CBS 623.88 were obtained from CBS-KNAW Collections (www.cbs.knaw.nl). Fungi were pre-grown on malt-extract agar (MEA, containing 20 g malt extract (Oxoid) and 12 g agar per liter). Inhibition experiments were performed on two types of agar medium, WYA5m and MEA. Methanol was also added to MEA for all *Methylovirgula* strains. Bacteria were inoculated onto the plates and incubated at 20 °C for 3 weeks for the *Methylovirgula* strains, and for 1 week for the other strains. In order to detect the production of antifungal compounds, agar plugs from *S. sanguinolentum* or *M. galericulata* cultures were inoculated in the middle of the plate at a distance of 2 cm from the pre-inoculated bacterial colonies. Plates were further incubated at 20 °C and inhibition zones or delayed fungal growth was observed during two months. For the detection of antibacterial compounds, bacteria were grown on WYA5m and MAE and overlay assays with *E. coli* WA123, *S. aureus* 533R4, *Dyella* WH32 and *Burkholderia* 4-A6 were performed as described above. Three replicates were prepared in all experiments, and delayed or inhibited fungal growth should be observed in all replicates.

## Results

In order to get an overview of the abundance and identity of bacteria that can be isolated from decaying wood, different sources of wood were used: 1) birch wood with either *Piptoporus betulinus*, *Fomes fomentarius*, *Stereum subtomentosum* or *Plicaturopsis crispa* fruiting bodies, 2) Abies fir and pine wood with clear visual indications of brown-rotting patterns, and 3) oak stumps several years after cutting. The number of bacteria varied between 0 and 1*10^8^ CFU per gram dry wood, with extremely low numbers in the birch wood samples (Figure 1a, Suppl. Table 2). The detection limit for the birch wood samples was about 200 CFU per gram dry wood. Low bacterial abundance in *Piptoporus* and *Fomes* colonized birch wood was confirmed on several independent sampling occasions (data not shown). Bacteria from Abies, pine and oak were isolated and identified by nearly complete 16S rDNA sequences (Suppl. Table 3). An average of 54% of randomly picked isolates had less than 97% sequence identity to known bacterial species, indicating that they were potentially new species. Growth of the *Methylovirgula* and *Beijerinckia* isolates slowed down after transfer to fresh agar plates without methanol and resumed after the addition of 1 g L^−1^ methanol to the growth medium.

**Figure 1:**
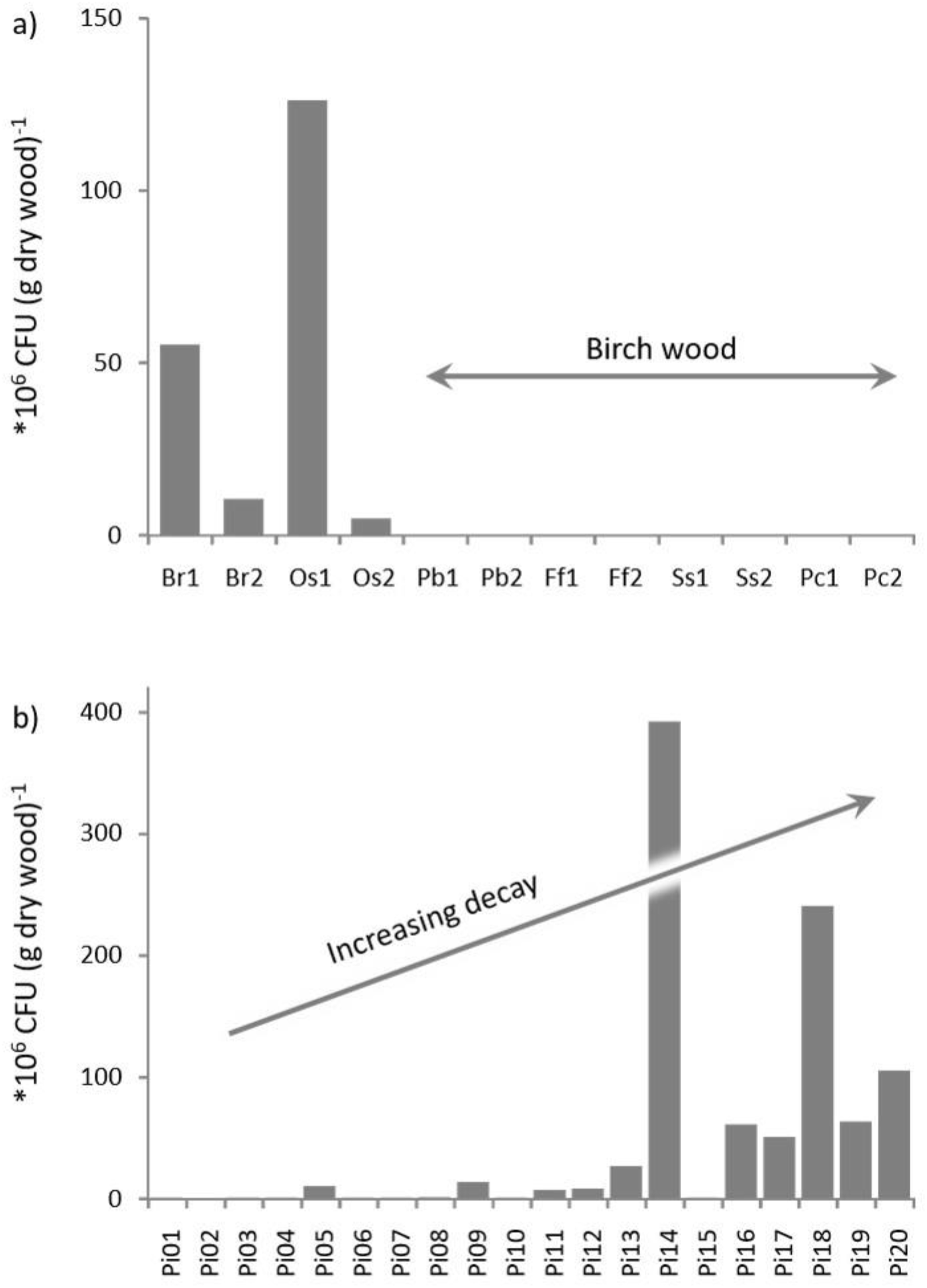
Number of colony forming units in decaying wood samples. a) Pilot experiment with decaying Abies (Br1), pine (Br2), oak (Os1, Os2) and birch wood (Pb1, Pb2, Ff1, Ff2, Ss1, Ss2, Pc1, Pc2). b) Main experiment with a decay range of pine wood. For a description of wood samples see Table 1 and Suppl. Table 2.

Bacterial communities were subsequently characterized in pine wood samples in different stages of decay. The number of bacteria, as determined by the number of CFU per gram dry weight, increased with decreasing wood density, which is an indicator for wood decay stage (Figure 1b, Table 1). Apart from density, wood moisture content and C/N ratio were also highly correlated with bacterial numbers (Suppl. Table 4). This was expected, because moisture increases, while C/N ratio decreases during decay. Moisture was used as a proxy for decay stage in a comparison of bacterial abundance in the experiments in this publication and the data of Valaskova et al (2009) (Figure 2). Other parameters that were measured, i.e. pH, ergosterol, manganese peroxidase, laccase and cellulose activity, did not correlate with the number of bacteria (Suppl. Table 4).

**Table 1:** Overview of wood parameters and number of bacteria.

| Wood sample | pH | Moisture (w/w) | Density (g (cm<sup>3</sup>)<sup>-1</sup>) | Nitrogen | Carbon | C/N | Ergosterol (mg (kg dry wood)<sup>-1</sup>) | CFU after 10 days (CFU (g dry wood)<sup>-1</sup>) | CFU after 3 weeks (CFU (g dry wood)<sup>-1</sup>) | CFU after 6 weeks (CFU (g dry wood)<sup>-1</sup>) | Mn-peroxidase* | Laccase* | Cellulase* |
| --- | --- | --- | --- | --- | --- | --- | --- | --- | --- | --- | --- | --- | --- |
| Pi01 | 5.1 | 22% | 0.50 | 0.07% | 49% | 674 | 12 | 5.0E+04 | 6.3E+04 | 6.3E+04 | 185 | 27 | 0 |
| Pi02 | 4.0 | 19% | 0.49 | 0.05% | 47% | 963 | 37 | 0.0E+00 | 0.0E+00 | 0.0E+00 | 15 | 0 | 0 |
| Pi03 | 4.3 | 19% | 0.45 | 0.02% | 51% | 2384 | 1 | 1.2E+04 | 1.2E+04 | 1.2E+04 | 6 | 0 | 0 |
| Pi04 | 4.9 | 28% | 0.42 | 0.05% | 47% | 911 | 14 | 2.6E+05 | 3.1E+05 | 3.1E+05 | 807 | 16 | 2 |
| Pi05 | 4.7 | 50% | 0.42 | 0.03% | 47% | 1421 | 17 | 9.7E+06 | 9.9E+06 | 9.9E+06 | 57 | 0 | 0 |
| Pi06 | 4.1 | 21% | 0.39 | 0.07% | 48% | 653 | 60 | 8.7E+04 | 1.6E+05 | 1.6E+05 | 22 | 0 | 3 |
| Pi07 | 3.7 | 25% | 0.37 | 0.05% | 48% | 1028 | 55 | 3.9E+04 | 5.1E+04 | 5.1E+04 | 191 | 0 | 1 |
| Pi08 | 4.2 | 22% | 0.37 | 0.07% | 47% | 653 | 29 | 8.9E+04 | 3.9E+05 | 5.2E+05 | 80 | 1 | 12 |
| Pi09 | 4.0 | 28% | 0.36 | 0.06% | 47% | 756 | 62 | 1.1E+07 | 1.2E+07 | 1.3E+07 | 7 | 0 | 1 |
| Pi10 | 4.2 | 25% | 0.35 | 0.06% | 47% | 734 | 28 | 5.1E+04 | 5.1E+04 | 6.4E+04 | 895 | 0 | 19 |
| Pi11 | 3.7 | 74% | 0.34 | 0.07% | 48% | 678 | 49 | 5.3E+05 | 3.8E+06 | 6.5E+06 | 108 | 1 | 2 |
| Pi12 | 3.8 | 76% | 0.32 | 0.07% | 47% | 665 | 28 | 7.4E+05 | 7.0E+06 | 7.9E+06 | 49 | 0 | 16 |
| Pi13 | 4.0 | 131% | 0.28 | 0.11% | 48% | 431 | 28 | 4.1E+06 | 1.6E+07 | 2.6E+07 | 101 | 0 | 0 |
| Pi14 | 4.0 | 146% | 0.23 | 0.22% | 52% | 243 | 44 | 4.6E+07 | 2.0E+08 | 3.9E+08 | 17 | 0 | 0 |
| Pi15 | 3.7 | 58% | 0.22 | 0.02% | 46% | 2143 | 51 | 5.7E+04 | 5.7E+04 | 5.7E+04 | 5 | 0 | 5 |
| Pi16 | 3.9 | 60% | 0.19 | 0.10% | 47% | 481 | 57 | 5.1E+07 | 6.1E+07 | 6.1E+07 | 55 | 0 | 2 |
| Pi17 | 4.4 | 128% | 0.19 | 0.44% | 48% | 108 | 207 | 6.6E+06 | 3.1E+07 | 5.0E+07 | 314 | 0 | 2 |
| Pi18 | 4.6 | 251% | 0.18 | 0.19% | 44% | 230 | 13 | 8.7E+07 | 1.0E+08 | 2.4E+08 | 48 | 0 | 0 |
| Pi19 | 4.8 | 237% | 0.15 | 0.41% | 50% | 125 | 57 | 1.0E+07 | 3.2E+07 | 6.3E+07 | 25 | 0 | 0 |
| Pi20 | 3.9 | 274% | 0.12 | 0.66% | 50% | 75 | 37 | 4.7E+07 | 7.7E+07 | 1.0E+08 | 76 | 0 | 0 |

\* Mn-peroxidase, laccase and cellulase activity in nmol DMAB-MBTH (g dry wood)<sup>-1</sup> h<sup>-1</sup>, μmol ABTS · (g dry wood)<sup>-1</sup> h<sup>-1</sup> and μmol RBB (g dry wood)<sup>-1</sup> h<sup>-1</sup> respectively.

**Figure 2:**
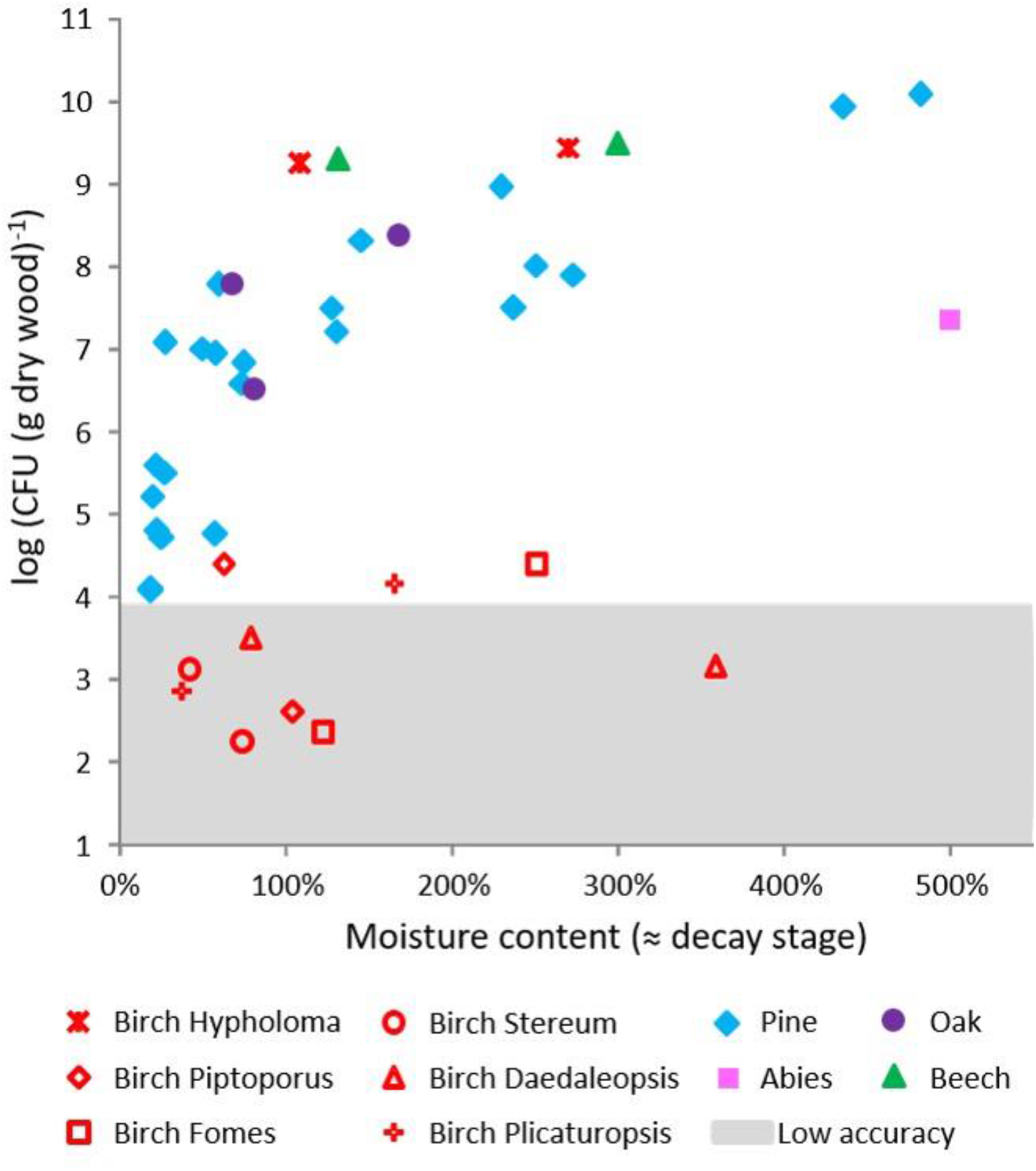
Number of colony forming units in decaying wood samples from the samples collected for this study and from Valášková *et al* (2009) combined. Moisture content serves as a proxy for decay stage, as density was not available for all samples. Measurements below 10^4^ CFU (g dry weight)^−1^ become less accurate due to the low number of colonies on plates, this area is indicated in grey.

Partial 16S rRNA sequences were determined for a maximum of 24 isolates per sample per time point. A complete list of identified bacteria is presented in Suppl. Table 5. Relative abundances of bacterial taxa were calculated for 14 of the 20 samples (Figure 3, Suppl. Table 6). For the other six samples, Pi01, Pi02, Pi03, Pi07, Pi10 and Pi15, only 0-3 sequences per sample were obtained. They were therefore excluded from the community analysis. Three genera were significantly correlated with density, namely *Streptacidiphilus*, *Mycobacterium* and *Rhodanobacter* (Kendall’s tau-b: P=0.006, 0.006 and 0.009 respectively). All three were exclusively found in more decayed wood samples with a density lower than 0.3 g (cm^3^)^−1^. The identity of isolated strains matched well with previously published pyrosequencing data on the same wood samples (Kielak et al 2016). We obtained representative isolates for all taxa (until genus level) with an abundance of more than 2% in the pyrosequencing data (Figure 4).

**Figure 3:**
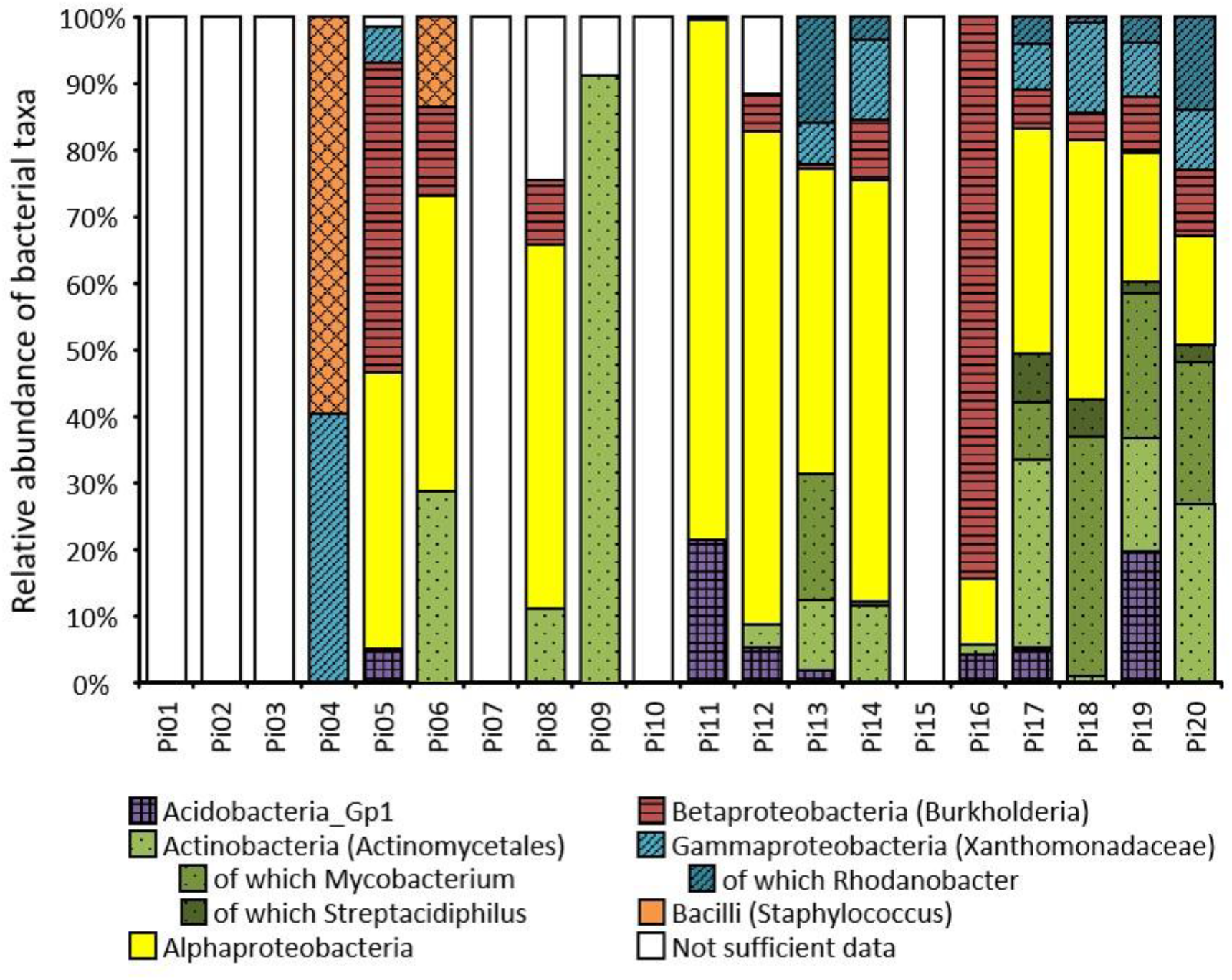
Bacterial community composition in decaying pine wood as determined by the identity of randomly picked isolates. For samples with very low bacterial abundance, we could not obtain sufficient data.

**Figure 4:**
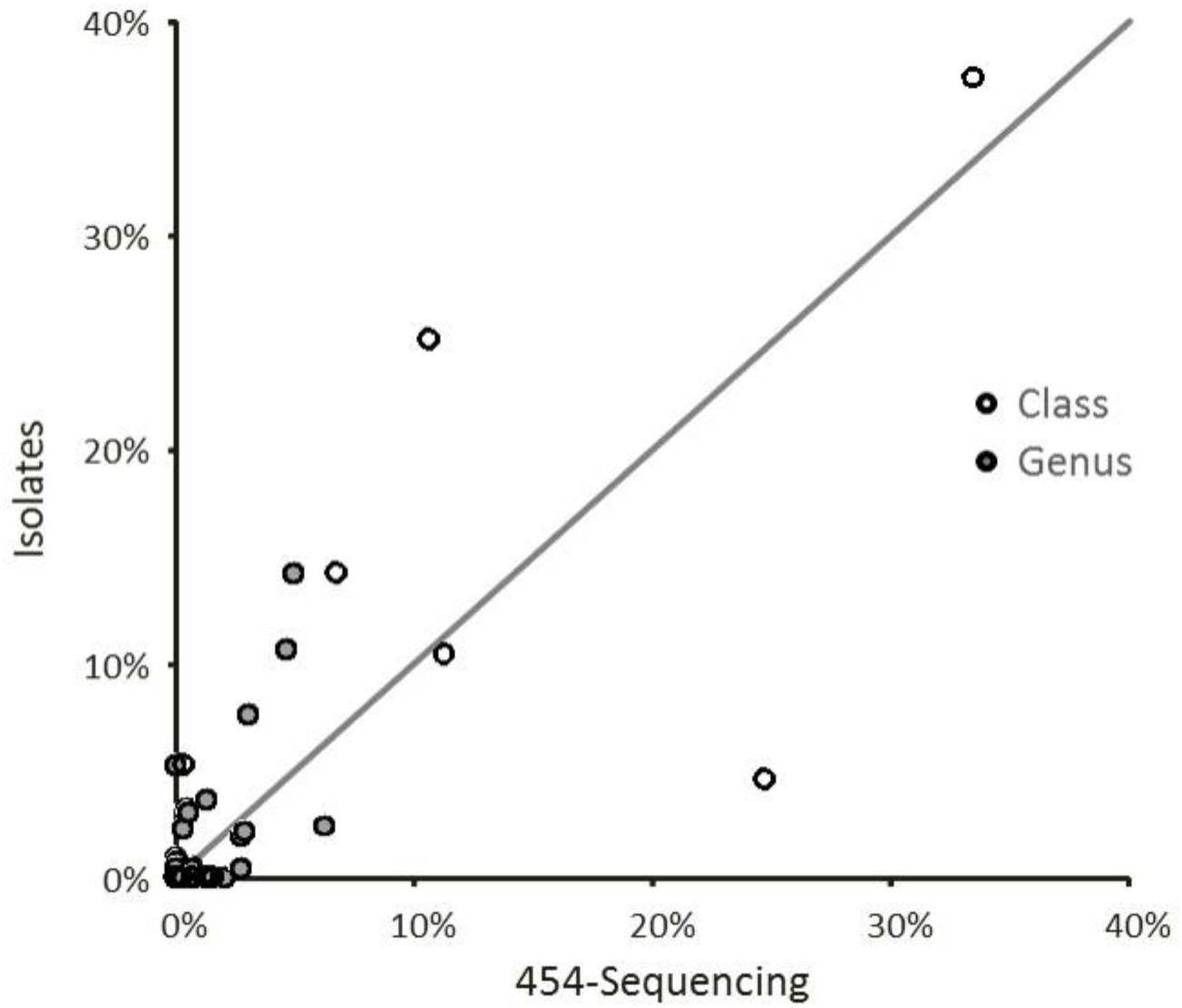
Relationship between the relative abundance of bacterial classes (open circles) and genera (closed circles) in pyrosequencing data and 16S rDNA sequences of isolated strains.

In order to identify those bacterial taxa that are most common in naturally decaying wood, we made a comparison between bacteria isolated in the current experiments and two published studies (Table 2). In the published work, bacteria were isolated from decaying wood of various tree species with the white-rot fungi *Hypholoma fascicularum* and *Resinicium bicolor* (Folman et al, 2008, Valaskova et al, 2009). Within the Alphaproteobacteria, members of the family *Acetobacteraceae* and the genus *Sphingomonas* were commonly isolated from decaying wood. *Methylovirgula*, one of the dominant taxa in decaying wood, was found in all studies except by Valaskova et al. Furthermore, *Burkholderia* sp. from the Betaproteobacteria and *Xanthomonadaceae* from the Gammaproteobacteria were recovered in every experiment in the comparison. Acidobacteria from subdivision 1 also appeared to be commonly isolated from decaying wood, with *Granulicella* as the most abundant genus.

**Table 2:**
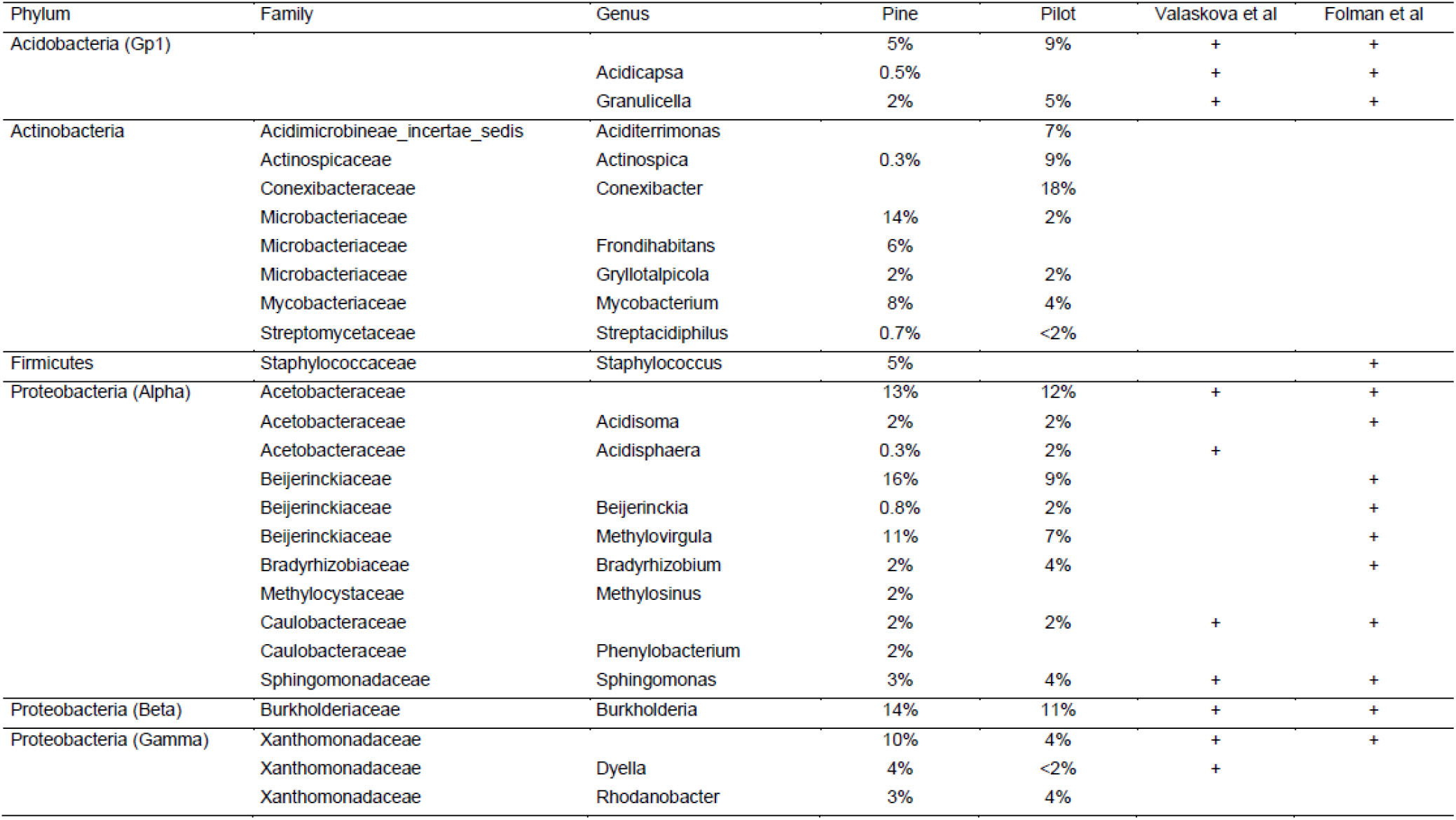
Comparison of bacterial taxa that were isolated from decaying wood in different experiments, the pilot and main experiment of this study, the experiment by Valášková et al (2009) and by Folman et al (2008). Strains that could not be identified at the family level were excluded, except for Acidobacteria. Taxa with only one or two representatives over all five experiments were also excluded.

Less than 2% of the isolates showed antibacterial activity. These bacteria were identified as *Streptacidiphilus* (5), *Kitasatospora* (1), Microbacteriaceae (1) and *Acidisoma* (3). They were active against *S. aureus* 533R4, *Dyella* WH32 and/or *Burkholderia* 4-A6. Antibacterial activity was observed in three wood samples, Pi16, Pi17 and Pi19, all of which belonged to the more decayed wood samples (Table 3). A selection of 36 strains of frequently isolated taxa were tested for antibacterial and antifungal activity on two different media (Table 4). The *Methylovirgula* strains did not grow on MEA and were therefore tested on WYA5m only. As in the first screening, antibacterial activity was not observed on the poor water-yeast agar, while two strains showed slight inhibition of *E. coli* on the rich MEA medium. In contrast, antifungal activity was observed against *S. sanguinolentum* and *M. galericulata* for several Xanthomonadaceae and *Burkholderia* strains. Three of the *Xanthomonadaceae* strains originally co-occurred with *S. sanguinolentum*, but had activity against both fungi. One *Mycobacterium* strain, isolated from late stage decay where it co-occurred with *M. galericulata* showed activity against *M. galericulata*, but not against *S. sanguinolentum* (Table 4). For *Burkholderia*, antifungal activity was more frequently directed against not co-occurring fungi than against co-occurring fungi.

**Table 3:** Isolates with antibacterial activity.

| Wood sample | Isolate | Selection | Taxon | Antibacterial activity against |
| --- | --- | --- | --- | --- |
| Pi16 | P26-C5 | random | Microbacteriaceae | Dyella |
| Pi16 | P26-B6 | random | Acidisoma | Dyella, Burkholderia |
| Pi16 | P26-C2 | random | Acidisoma | Dyella, Burkholderia |
| Pi16 | P26-B4 | random | Acidisoma | Dyella, Burkholderia |
| Pi17 | P43-C6 | random | Kitasatospora | Staphylococcus, Dyella |
| Pi17 | P03-D6 | random | Streptacidiphilus | Staphylococcus |
| Pi17 | P43-C3 | random | Streptacidiphilus | Staphylococcus, Dyella |
| Pi17 | P43-D4 | random | Streptacidiphilus | Staphylococcus, Dyella |
| Pi17 | P03-A5 | morphology | Streptacidiphilus | Staphylococcus, Dyella |
| Pi19 | P43-B6 | random | ?? | Staphylococcus |
| Pi19 | P02-A3 | morphology | Streptacidiphilus | Dyella |

\* No sequence could be obtained for this isolate.

**Table 4:**
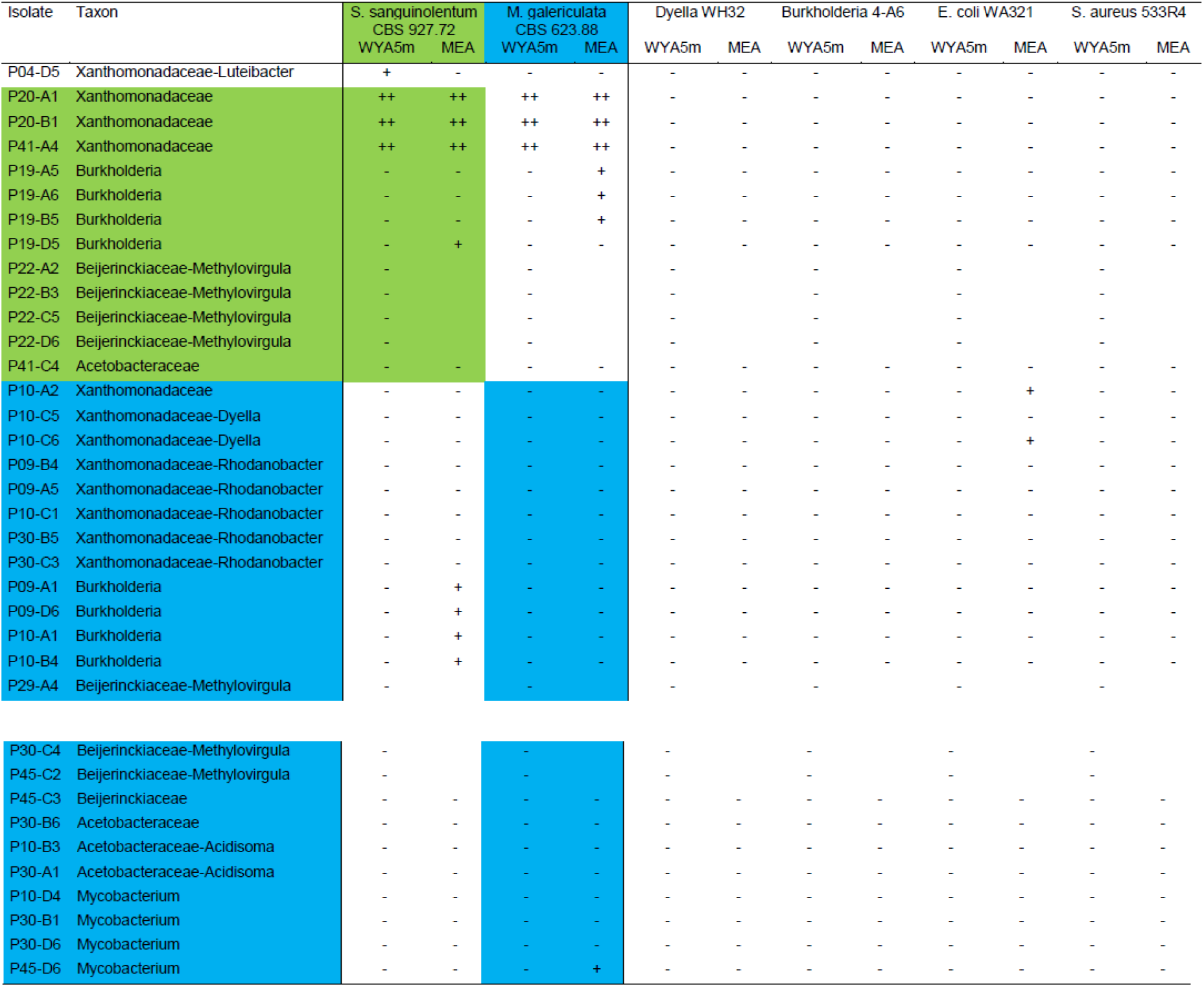
Antifungal and antibacterial activity of bacteria from decaying wood. Green and blue indicate bacteria that originally co-occurred with *S. sanguinolentum* and *M. galericulata* respectively. ++, inhibited fungal growth; +, delayed fungal growth; -, no effect on fungal growth.

## Discussion

Antagonistic interactions among wood decaying fungi have been extensively studied, but little is known about the interactions with other wood inhabiting microorganisms (Johnston *et al* 2016). We know that wood decaying fungi heavily compete for substrates and a wide range of antimicrobial compounds have been identified (Hiscox et al 2015). These fungal compounds can inhibit fungi as well as bacteria (Bérdy 2005). Toxic fungal compounds could potentially be an explanation for the extremely low abundance of bacteria in decaying birch wood. Though low bacterial numbers were observed for four different wood decay fungi, including white and brown-rot fungi from three different orders (*Polyporales*, *Russulales* and *Agricales*), high numbers have also been observed for birch wood colonized by *Hypholoma fasciculare* (Valášková et al 2009). Therefore, we can conclude that the abundance of bacteria in decaying birch wood is influenced by fungal species. Another pattern was observed for decaying pine wood. We showed that bacterial abundance was correlated with wood density, which is an indicator for the stage of wood decay. Later stages of decay were apparently more favorable for bacteria than earlier stages, which might be related to the higher moisture and nitrogen content. However, there is one exception in this range. Sample Pi15 was highly decayed, while bacterial numbers were around the limit of detection. This sample was dominated by one fungal species, 99.7% *Ischnoderma benzoinum*. To conclude, the general pattern is that bacterial abundance increases during wood decay. In addition, we identified several wood decaying fungi that successfully excluded other fungi and bacteria. These were *Piptoporus betulinus*, *Fomes fomentarius*, *Stereum subtomentosum*, *Plicaturopsis crispa* and *Ischnoderma benzoinum*.

In order to investigate antagonistic activity of bacteria from decaying wood against fungi and other bacteria, we established a culture collection of bacteria from decaying wood. One of the major problems when working with isolated bacteria is that they are often not representative for the original community. In most habitats, only a few percent of the bacteria is culturable and many taxa are not captured in isolation attempts (Rappe and Giovannoni 2003). Valášková et al (2009) previously demonstrated that it is relatively easy to culture bacteria from decaying wood, even for groups that are generally considered as hard-to-culture like the *Acidobacteria* (Jones et al 2009). In agreement, we found a good match between the identity of isolated bacteria and pyrosequencing data of the same wood samples (Kielak *et al* 2016). Though *Acidobacteria* were underrepresented in the culture collection, we obtained representative isolates for all taxa (until genus level) with an abundance of more than 2% in the pyrosequencing data. In addition, many potentially new bacterial species were isolated. Several new groups within the genus *Methylovirgula* were also captured. It is likely that these groups were previously missed, because of the absence of methanol in the isolation medium. By comparing our data with previously published studies reporting on isolated bacteria from decaying wood, we were able to identify several taxa that are ‘typical’ for this environment. These were *Xanthomonodaceae*, *Acetobacteraceae*, *Caulobacteraceae*, *Methylovirgula*, *Sphingomonas*, *Burkholderia* and *Granulicella*. Our current collection is highly representative for bacterial communities in decaying wood.

The complete collection of bacteria from decaying pine wood was screened for antagonistic activity against four different bacteria, but surprisingly few bacteria produced antagonistic compounds against other bacteria. Low bacterial abundance could be an explanation for low antibacterial activity, because bacteria are less likely to encounter each other at low abundance. Indeed, all bacteria that exhibited antibacterial activity were isolated from the more decayed wood samples with higher bacterial abundance. However, even in those samples, antibacterial activity was not very common. Screening for antibacterial activity was performed on a low nutrient agar, water-yeast agar pH5 with methanol. We choose this medium, because many isolates are not able to grow at higher nutrient concentrations or pH. It is likely that the medium composition influenced the production of antimicrobial compounds. Higher nutrient concentrations did not increase the frequency of bacteria expressing antibacterial compounds in a pilot experiment (data not shown), though a slight increase was observed when a subset of the isolates from pine wood were tested on MEA medium (Table 4). Apparently, bacterial competition via the production of antibacterial compounds is not an important trait for bacteria in wood decay.

We originally hypothesized that bacteria would not attack the wood decaying fungi, because they are thought to be nutritionally dependent on the fungi. However, in contrast to observations by Valášková et al (2009), several isolates from decaying wood showed antagonistic activity against wood decaying fungi. *Xanthomonadaceae* isolated from wood colonized by the early successional fungus *S. sanguinolentum* were antagonistic against *S. sanguinolentum* as well as the late successional fungus *M. galericulata*. For *Burkholderia*, the isolates from *S. sanguinolentum* colonized wood were pre-dominantly active against *M. galericulata* and *vice versa*. So, no general pattern could be observed related to decay stage or co-occurrence, but antifungal activity was definitely higher than antibacterial activity. Close associations between soil *Burkholderia* and fungi have been reported with antagonistic as well as mutualistic interactions (Stopnisek 2016). Members of the genera *Burkholderia* and *Dyella* (*Xanthomonadaceae*) have both been identified as mycophagous, *i.e*. fungal feeding bacteria (Rudnik et al 2015; Ballhausen et al 2015) and are able to migrate along fungal hyphae (Warmink and van Elsas 2009). The picture arises of bacteria that are consuming fungal hyphae and/or fungal exudates, while at the same time using the hyphae to explore new territories. Assuming that the bacteria are not able to totally eliminate the wood decaying fungi, this is likely to be an efficient strategy. Feeding on fungi has also been indicated as an important nutritional strategy of bacteria inhabiting decomposing leaf litter (Tláskal *et al*., 2016).

In conclusion, bacterial abundance in decaying wood increases with progressive wood decay, but can be drastically diminished when wood is colonized by some specific fungi. Contrary to our original hypothesis, bacteria isolated from decaying wood hardly exhibited antagonistic activity against bacteria, while several isolates did show antagonistic activity against wood decaying fungi.

## Acknowledgements

We thank Maria Hundscheid for technical assistance. Natuurmonumenten kindly provided access to the field site and gave permission to take samples. This is a NIOO publication no. xxxx.

