## supplementary tables for "Identification and antimicrobial properties of bacteria isolated from naturally decaying wood"

Supplementary Tables with:  Scheublin et al. Identification and antimicrobial properties of bacteria isolated from naturally decaying wood.

Suppl. Table 1: Selected bacterial isolates for in-depth characterization of antimicrobial properties.
The relative abundance of *Stereum sanguinolentum* and *Mycena galericulata* in the original pine wood sample was determined by pyrosequencing (Kielak et al 2016).

| Isolate | Family | Genus | Wood sample | Decay stage* | S. sanguinolentum | M. galericulata |
|---|---|---|---|---|---|---|
| P04-D5 | Xanthomonadaceae | Luteibacter | Pi05 | Early | 0% | 0% |
| P20-A1 | Xanthomonadaceae | unclassified | Pi04 | Early | 35% | 0% |
| P20-B1 | Xanthomonadaceae | unclassified | Pi04 | Early | 35% | 0% |
| P41-A4 | Xanthomonadaceae | unclassified | Pi04 | Early | 35% | 0% |
| P19-A5 | Burkholderiaceae | Burkholderia | Pi08 | Middle | 54% | 0% |
| P19-A6 | Burkholderiaceae | Burkholderia | Pi08 | Middle | 54% | 0% |
| P19-B5 | Burkholderiaceae | Burkholderia | Pi08 | Middle | 54% | 0% |
| P19-D5 | Burkholderiaceae | Burkholderia | Pi06 | Middle | 8% | 0% |
| P22-A2 | Beijerinckiaceae | Methylovirgula | Pi08 | Middle | 54% | 0% |
| P22-B3 | Beijerinckiaceae | Methylovirgula | Pi08 | Middle | 54% | 0% |
| P22-C5 | Beijerinckiaceae | Methylovirgula | Pi08 | Middle | 54% | 0% |
| P22-D6 | Beijerinckiaceae | Methylovirgula | Pi08 | Middle | 54% | 0% |
| P41-C4 | Acetobacteraceae | unclassified | Pi06 | Middle | 8% | 0% |
| P10-A2 | Xanthomonadaceae | unclassified | Pi13 | Late | 0% | 27% |
| P10-C5 | Xanthomonadaceae | Dyella | Pi13 | Late | 0% | 27% |
| P10-C6 | Xanthomonadaceae | Dyella | Pi13 | Late | 0% | 27% |
| P09-B4 | Xanthomonadaceae | Rhodanobacter | Pi20 | Late | 0% | 23% |
| P09-A5 | Xanthomonadaceae | Rhodanobacter | Pi20 | Late | 0% | 23% |
| P10-C1 | Xanthomonadaceae | Rhodanobacter | Pi13 | Late | 0% | 27% |
| P30-B5 | Xanthomonadaceae | Rhodanobacter | Pi13 | Late | 0% | 27% |
| P30-C3 | Xanthomonadaceae | Rhodanobacter | Pi13 | Late | 0% | 27% |
| P09-A1 | Burkholderiaceae | Burkholderia | Pi20 | Late | 0% | 23% |
| P09-C3 | Burkholderiaceae | Burkholderia | Pi20 | Late | 0% | 23% |
| P10-A1 | Burkholderiaceae | Burkholderia | Pi13 | Late | 0% | 27% |
| P10-B4 | Burkholderiaceae | Burkholderia | Pi13 | Late | 0% | 27% |
| P29-A4 | Beijerinckiaceae | Methylovirgula | Pi20 | Late | 0% | 23% |
| P30-C4 | Beijerinckiaceae | Methylovirgula | Pi13 | Late | 0% | 27% |
| P45-C2 | Beijerinckiaceae | Methylovirgula | Pi13 | Late | 0% | 27% |

Suppl. Table 2: Description of decaying wood samples in pilot experiment.

| Sample | Description | pH | Moisture (w/w) | CFU per gram dry wood |
|---|---|---|---|---|
| Pb1 | Birch with *Piptoporus betulinus* fruiting body | 3,30 | 105% | 4,1E+02 |
| Pb2 | Birch with *Piptoporus betulinus* fruiting body | 3,41 | 64% | 2,4E+04 |
| Ff1 | Birch with *Fomes fomentarius* fruiting body | 4,88 | 252% | 2,4E+04 |
| Ff2 | Birch with *Fomes fomentarius* fruiting body | 4,25 | 124% | 0,0E+00 |
| Ss1 | Birch with *Stereum subtomentosum* fruiting body | 3,51 | 75% | 1,8E+02 |
| Ss2 | Birch with *Stereum subtomentosum* fruiting body | 4,02 | 43% | 1,3E+03 |
| Pc1 | Birch with *Plicaturopsis crispa* fruiting body | 3,74 | 39% | 6,9E+02 |
| Pc2 | Birch with *Plicaturopsis crispa* fruiting body | 4,41 | 167% | 1,4E+04 |
| Br1 | Abies fir with brown-rot | 4,35 | 500% | 5,5E+07 |
| Br2 | Pine with brown-rot | 4,77 | 58% | 1,1E+07 |
| Os1 | Oak stump 2 years after cutting | n.d. | 68% | 1,3E+08 |
| Os2 | Oak stump 5 years after cutting | n.d. | 82% | 5,0E+06 |

n.d. = not determined

Suppl. Table 3: Identification of bacteria isolated from decaying wood samples in pilot experiment.
Identification was based on nearly complete 16S rRNA gene sequences.
Sequences with less than 97% sequence identify to known bacterial species are indicated in bold.

| Sample | Isolate | Selection | Identification by EzTaxon | | | Classification by RDP Classifier |
|---|---|---|---|---|---|---|
| | | | Closest match | Similarity | Completeness | |
| Br1 | Br1-1 | random | Granulicella paludicola OB1010(T) | 99 | 95 | Granulicella sp. |
| Br1 | Br1-2 | random | Granulicella paludicola OB1010(T) | 99 | 94 | Granulicella sp. |
| Br1 | Br1-3 | random | Granulicella arctica MP5ACTX2(T) | 97 | 94 | Granulicella sp. |
| Br1 | Br1-4 | random | Reyranella massiliensis 521(T) | 99 | 91 | Alphaproteobacteria |
| **Br1** | **Br1-5** | **random** | **Kozakia baliensis Yo-3(T)** | **95** | **95** | **Acetobacteraceae** |
| **Br1** | **Br1-6** | **random** | **'Rhodovastum atsumiense' G2-11(T)** | **97** | **92** | **Acidisphaera sp.** |
| Br1 | Br1-7 | random | Methylovirgula ligni BW863(T) | 99 | 95 | Methylovirgula sp. |
| Br1 | Br1-8 | random | Methylovirgula ligni BW863(T) | 99 | 93 | Methylovirgula sp. |
| Br1 | Br1-9 | random | Methylovirgula ligni BW863(T) | 99 | 94 | Methylovirgula sp. |
| Br1 | Br1-10 | random | Bradyrhizobium lablabi CCBAU 23086(T) | 99 | 95 | Bradyrhizobium sp. |
| Br1 | Br1-11 | random | Burkholderia sartisoli RP007(T) | 98 | 95 | Burkholderia sp. |
| Br1 | Br1-12 | random | Burkholderia sordidicola S5-B(T) | 98 | 96 | Burkholderia sp. |
| Br1 | Br1-13 | random | Rhodanobacter umsongensis GR24-2(T) | 98 | 93 | Rhodanobacter sp. |
| Br1 | Br1-14 | not random | Mycobacterium phocaicum CIP 108542(T) | 98 | 92 | Mycobacterium sp. |
| Br1 | Br1-15 | not random | Mycobacterium llatzerense MG13(T) | 98 | 95 | Mycobacterium sp. |
| Br1 | Br1-16 | not random | Rhodococcus globerulus DSM 4954(T) | 99 | 95 | Rhodococcus sp. |
| Br1 | Br1-17 | not random | Streptacidiphilus neutrinimicus DSM 41755(T) | 100 | 93 | Streptacidiphilus sp. |
| **Br1** | **Br1-18** | **not random** | **Kozakia baliensis Yo-3(T)** | **95** | **94** | **Acetobacteraceae** |
| **Br1** | **Br1-19** | **not random** | **Kozakia baliensis Yo-3(T)** | **95** | **94** | **Acetobacteraceae** |
| Br1 | Br1-20 | not random | Methylovirgula ligni BW863(T) | 99 | 94 | Methylovirgula sp. |
| Br1 | Br1-21 | not random | Bradyrhizobium japonicum USDA 6(T) | 99 | 95 | Bradyrhizobium sp. |
| Br1 | Br1-22 | not random | Bradyrhizobium huanghuaihaiense CCBAU23303(T) | 99 | 95 | Bradyrhizobium sp. |
| Br1 | Br1-23 | not random | Burkholderia phenazinium LMG 2247(T) | 99 | 96 | Burkholderia sp. |
| Br1 | Br1-24 | not random | Dyella soli JS12-10(T) | 99 | 95 | Dyella sp. |
| **Br1** | **Br1-25** | **not random** | **Steroidobacter denitrificans FS(T)** | **90** | **94** | **Steroidobacter sp.** |
| **Br2** | **Br2-1** | **random** | **Conexibacter arvalis KV-962(T)** | **96** | **92** | **Conexibacter sp.** |
| **Br2** | **Br2-2** | **random** | **Conexibacter arvalis KV-962(T)** | **96** | **91** | **Conexibacter sp.** |

Suppl. Table 4: Correlation matrices ( Kendall's Tau b and Spearman's rho) between wood parameters and the number of cultivable bacteria (CFU)

| | | Density | Moisture | pH | C/N | Ergosterol | MnPeroxidase | Laccase | Cellulase | CFU/g dry |
|---|---|---|---|---|---|---|---|---|---|---|
| **Kendall** | | | | | | | | | | |
| Density | Correlation Coefficient | 1,000 | -.705** | ,179 | .589** | -,305 | ,053 | .405* | -,043 | -.611** |
| | Sig. (2-tailed) | | ,000 | ,281 | ,000 | ,060 | ,746 | ,026 | ,794 | ,000 |
| | N | 20 | 20 | 20 | 20 | 20 | 20 | 20 | 20 | 20 |
| Moisture | Correlation Coefficient | -.705** | 1,000 | -,060 | -.484** | ,095 | ,032 | -,258 | -,140 | .674** |
| | Sig. (2-tailed) | ,000 | | ,719 | ,003 | ,559 | ,846 | ,158 | ,396 | ,000 |
| | N | 20 | 20 | 20 | 20 | 20 | 20 | 20 | 20 | 20 |
| pH | Correlation Coefficient | ,179 | -,060 | 1,000 | -,103 | -,298 | ,136 | ,349 | -,327 | ,049 |
| | Sig. (2-tailed) | ,281 | ,719 | | ,535 | ,072 | ,414 | ,062 | ,052 | ,769 |
| | N | 20 | 20 | 20 | 20 | 20 | 20 | 20 | 20 | 20 |
| C/N | Correlation Coefficient | .589** | -.484** | -,103 | 1,000 | -,189 | -,105 | ,169 | ,065 | -.600** |
| | Sig. (2-tailed) | ,000 | ,003 | ,535 | | ,243 | ,516 | ,353 | ,695 | ,000 |
| | N | 20 | 20 | 20 | 20 | 20 | 20 | 20 | 20 | 20 |
| Ergosterol | Correlation Coefficient | -,305 | ,095 | -,298 | -,189 | 1,000 | -,116 | -.376* | ,204 | ,147 |
| | Sig. (2-tailed) | ,060 | ,559 | ,072 | ,243 | | ,475 | ,040 | ,215 | ,364 |
| | N | 20 | 20 | 20 | 20 | 20 | 20 | 20 | 20 | 20 |
| MnPeroxidase | Correlation Coefficient | ,053 | ,032 | ,136 | -,105 | -,116 | 1,000 | .479** | ,172 | -,063 |
| | Sig. (2-tailed) | ,746 | ,846 | ,414 | ,516 | ,475 | | ,009 | ,296 | ,697 |
| | N | 20 | 20 | 20 | 20 | 20 | 20 | 20 | 20 | 20 |
| Laccase | Correlation Coefficient | .405* | -,258 | ,349 | ,169 | -.376* | .479** | 1,000 | ,083 | -,243 |
| | Sig. (2-tailed) | ,026 | ,158 | ,062 | ,353 | ,040 | ,009 | | ,656 | ,183 |
| | N | 20 | 20 | 20 | 20 | 20 | 20 | 20 | 20 | 20 |
| Cellulase | Correlation Coefficient | -,043 | -,140 | -,327 | ,065 | ,204 | ,172 | ,083 | 1,000 | -,183 |
| | Sig. (2-tailed) | ,794 | ,396 | ,052 | ,695 | ,215 | ,296 | ,656 | | ,267 |
| | N | 20 | 20 | 20 | 20 | 20 | 20 | 20 | 20 | 20 |
| CFU/g dry | Correlation Coefficient | -.611** | .674** | ,049 | -.600** | ,147 | -,063 | -,243 | -,183 | 1,000 |
| | Sig. (2-tailed) | ,000 | ,000 | ,769 | ,000 | ,364 | ,697 | ,183 | ,267 | |
| | N | 20 | 20 | 20 | 20 | 20 | 20 | 20 | 20 | 20 |

Suppl. Table 5: Complete list of bacteria isolated from decaying pine wood samples

| Wood sample | Isolate | Time | Selection | Antibacterial activity | Length [bp] | Identification by EzTaxon<br>Closest match | Similarity | Completeness |
|---|---|---|---|---|---|---|---|---|
| Pi04 | P20-A1 | 10 days | random | No | 988 | Dyella ginsengisoli Gsoil 3046(T) | 99 | 67 |
| Pi04 | P20-A4 | 10 days | random | No | 910 | Staphylococcus devriesei LMG 25332(T) | 91 | 61 |
| Pi04 | P20-B1 | 10 days | random | No | 689 | Frateuria aurantia DSM 6220(T) | 99 | 47 |
| Pi04 | P20-C2 | 10 days | random | No | 803 | Staphylococcus warneri ATCC 27836(T) | 100 | 55 |
| Pi04 | P20-C3 | 10 days | random | No | 797 | Staphylococcus warneri ATCC 27836(T) | 100 | 54 |
| Pi04 | P20-C4 | 10 days | random | No | 653 | Staphylococcus warneri ATCC 27836(T) | 100 | 44 |
| Pi04 | P20-D1 | 10 days | random | No | 781 | Staphylococcus warneri ATCC 27836(T) | 100 | 53 |
| Pi04 | P41-A4 | 3 weeks | random | No | 982 | Dyella ginsengisoli Gsoil 3046(T) | 99 | 67 |
| Pi05 | P04-A1 | 10 days | morphology | No | 862 | Luteibacter rhizovicinus LJ96(T) | 100 | 59 |
| Pi05 | P04-A2 | 10 days | morphology | No | 839 | Luteibacter rhizovicinus LJ96(T) | 100 | 57 |
| Pi05 | P04-A3 | 10 days | random | No | 862 | Burkholderia oklahomensis C6786(T) | 98 | 59 |
| Pi05 | P04-A4 | 10 days | random | No | 723 | Burkholderia stabilis LMG 14294(T) | 98 | 50 |
| Pi05 | P04-A5 | 10 days | random | No | 853 | Burkholderia oklahomensis C6786(T) | 98 | 59 |
| Pi05 | P04-A6 | 10 days | random | No | 878 | 'Sphingomonas kyeonggiense' THG-DT81(T) | 96 | 62 |
| Pi05 | P04-B1 | 10 days | random | No | 883 | Burkholderia oklahomensis C6786(T) | 98 | 61 |
| Pi05 | P04-B2 | 10 days | random | No | 837 | Burkholderia oklahomensis C6786(T) | 98 | 58 |
| Pi05 | P04-B3 | 10 days | random | No | 822 | Sphingomonas polyaromaticivorans B2-7(T) | 98 | 58 |
| Pi05 | P04-B4 | 10 days | random | No | 867 | Sphingomonas oligoaromativorans SY-6(T) | 99 | 62 |
| Pi05 | P04-B5 | 10 days | random | No | 772 | Sphingomonas polyaromaticivorans B2-7(T) | 98 | 55 |
| Pi05 | P04-B6 | 10 days | random | No | 852 | Burkholderia oklahomensis C6786(T) | 98 | 59 |
| Pi05 | P04-C1 | 10 days | random | No | 867 | Burkholderia oklahomensis C6786(T) | 98 | 60 |
| Pi05 | P04-C2 | 10 days | random | No | 858 | Granulibacter bethesdensis CGDNIH1(T) | 95 | 61 |
| Pi05 | P04-C4 | 10 days | random | No | 505 | Sphingomonas polyaromaticivorans B2-7(T) | 99 | 36 |
| Pi05 | P04-C5 | 10 days | random | No | 864 | Terriglobus roseus DSM 18391(T) | 98 | 61 |
| Pi05 | P04-C6 | 10 days | random | No | 857 | Sphingomonas dokdonensis DS-4(T) | 97 | 61 |
| Pi05 | P04-D1 | 10 days | random | No | 841 | Burkholderia sordidicola S5-B(T) | 100 | 58 |
| Pi05 | P04-D3 | 10 days | random | No | 830 | Burkholderia oklahomensis C6786(T) | 98 | 57 |
| Pi05 | P04-D5 | 10 days | random | No | 837 | Luteibacter rhizovicinus LJ96(T) | 100 | 57 |
| Pi05 | P04-D6 | 10 days | random | No | 846 | Sphingomonas dokdonensis DS-4(T) | 96 | 60 |

| | | | | | | | | |
|---|---|---|---|---|---|---|---|---|
| Pi06 | P19-C6 | 10 days | random | No | 1029 | Staphylococcus warneri ATCC 27836(T) | 100 | 70 |
| Pi06 | P19-D1 | 10 days | random | No | 854 | Phenylobacterium composti 4T-6(T) | 96 | 61 |
| Pi06 | P19-D2 | 10 days | random | No | 913 | Gryllotalpicola daejeonensis RU-04(T) | 99 | 63 |
| Pi06 | P19-D5 | 10 days | random | No | 852 | Burkholderia udeis LMG 27134(T) | 99 | 59 |
| Pi06 | P41-C3 | 3 weeks | random | No | 902 | Methylosinus trichosporium OB3b(T) | 95 | 65 |
| Pi06 | P41-C4 | 3 weeks | random | No | 1135 | Acidisphaera rubrifaciens HS-AP3(T) | 95 | 80 |
| Pi06 | P41-C6 | 3 weeks | random | No | 1031 | Gryllotalpicola ginsengisoli DSM 22003(T) | 98 | 71 |
| Pi07 | P19-C3 | 10 days | random | No | 819 | Gluconacetobacter tumulisoli T611xx-1-4a(T) | 97 | 58 |
| Pi07 | P19-C4 | 10 days | random | No | 844 | Gluconacetobacter tumulisoli T611xx-1-4a(T) | 97 | 59 |
| Pi07 | P19-C5 | 10 days | random | No | 905 | Gluconacetobacter tumulisoli T611xx-1-4a(T) | 97 | 64 |
| Pi08 | P19-A5 | 10 days | random | No | 892 | Burkholderia ambifaria AMMD(T) | 98 | 61 |
| Pi08 | P19-A6 | 10 days | random | No | 909 | Burkholderia ambifaria AMMD(T) | 98 | 62 |
| Pi08 | P19-B1 | 10 days | random | No | 865 | Sphingomonas polyaromaticivorans B2-7(T) | 99 | 61 |
| Pi08 | P19-B2 | 10 days | random | No | 774 | Sphingomonas oligoaromativorans SY-6(T) | 98 | 55 |
| Pi08 | P19-B3 | 10 days | random | No | 862 | Burkholderia ambifaria AMMD(T) | 98 | 59 |
| Pi08 | P19-B4 | 10 days | random | No | 896 | Sphingomonas oligoaromativorans SY-6(T) | 98 | 64 |
| Pi08 | P19-B5 | 10 days | random | No | 780 | Burkholderia stabilis LMG 14294(T) | 98 | 54 |
| Pi08 | P22-A2 | 3 weeks | random | No | 840 | Methylovirgula ligni BW863(T) | 99 | 60 |
| Pi08 | P22-A3 | 3 weeks | random | No | 791 | Methylovirgula ligni BW863(T) | 98 | 56 |
| Pi08 | P22-A4 | 3 weeks | random | No | 940 | Gryllotalpicola ginsengisoli DSM 22003(T) | 99 | 65 |
| Pi08 | P22-A5 | 3 weeks | random | No | 921 | Gryllotalpicola ginsengisoli DSM 22003(T) | 99 | 64 |
| Pi08 | P22-A6 | 3 weeks | random | No | 926 | Methylovirgula ligni BW863(T) | 99 | 66 |
| Pi08 | P22-B2 | 3 weeks | random | No | 932 | Methylovirgula ligni BW863(T) | 98 | 66 |
| Pi08 | P22-B3 | 3 weeks | random | No | 935 | Methylovirgula ligni BW863(T) | 99 | 67 |
| Pi08 | P22-B4 | 3 weeks | random | No | 926 | Methylovirgula ligni BW863(T) | 99 | 66 |
| Pi08 | P22-B5 | 3 weeks | random | No | 930 | Methylovirgula ligni BW863(T) | 99 | 66 |
| Pi08 | P22-B6 | 3 weeks | random | No | 925 | Methylovirgula ligni BW863(T) | 99 | 66 |
| Pi08 | P22-C1 | 3 weeks | random | No | 880 | Methylovirgula ligni BW863(T) | 99 | 63 |
| Pi08 | P22-C2 | 3 weeks | random | No | 870 | Methylovirgula ligni BW863(T) | 99 | 62 |
| Pi08 | P22-C3 | 3 weeks | random | No | 288 | Beijerinckia doebereinerae CECT 7311(T) | 98 | 20 |
| Pi08 | P22-C4 | 3 weeks | random | No | 872 | Methylovirgula ligni BW863(T) | 99 | 62 |
| Pi08 | P22-C5 | 3 weeks | random | No | 940 | Methylovirgula ligni BW863(T) | 99 | 67 |
| Pi08 | P22-D1 | 3 weeks | random | No | 803 | Methylovirgula ligni BW863(T) | 99 | 57 |

| | | | | | | | | |
|---|---|---|---|---|---|---|---|---|
| Pi08 | P22-D2 | 3 weeks | random | No | 920 | Gryllotalpicola ginsengisoli DSM 22003(T) | 99 | 64 |
| Pi08 | P22-D3 | 3 weeks | random | No | 975 | Gryllotalpicola ginsengisoli DSM 22003(T) | 99 | 68 |
| Pi08 | P22-D4 | 3 weeks | random | No | 877 | Methylovirgula ligni BW863(T) | 99 | 62 |
| Pi08 | P22-D5 | 3 weeks | random | No | 894 | Methylovirgula ligni BW863(T) | 99 | 64 |
| Pi08 | P22-D6 | 3 weeks | random | No | 937 | Methylovirgula ligni BW863(T) | 99 | 67 |
| Pi09 | P07-A1 | 10 days | morphology | No | 815 | Burkholderia udeis LMG 27134(T) | 98 | 56 |
| Pi09 | P07-A2 | 10 days | morphology | No | 754 | Frondihabitans peucedani RS-15(T) | 97 | 54 |
| Pi09 | P07-A3 | 10 days | morphology | No | 769 | Frondihabitans cladoniiphilus CafT13(T) | 100 | 53 |
| Pi09 | P07-A4 | 10 days | random | No | 746 | Frondihabitans peucedani RS-15(T) | 99 | 51 |
| Pi09 | P07-A5 | 10 days | random | No | 948 | Frondihabitans peucedani RS-15(T) | 100 | 65 |
| Pi09 | P07-A6 | 10 days | random | No | 767 | Frondihabitans cladoniiphilus CafT13(T) | 100 | 53 |
| Pi09 | P07-B1 | 10 days | random | No | 936 | Frondihabitans peucedani RS-15(T) | 100 | 65 |
| Pi09 | P07-B2 | 10 days | random | No | 923 | Frondihabitans peucedani RS-15(T) | 100 | 64 |
| Pi09 | P07-B3 | 10 days | random | No | 831 | Frondihabitans peucedani RS-15(T) | 100 | 57 |
| Pi09 | P07-B4 | 10 days | random | No | 578 | Frondihabitans cladoniiphilus CafT13(T) | 99 | 40 |
| Pi09 | P07-B5 | 10 days | random | No | 491 | Frondihabitans cladoniiphilus CafT13(T) | 100 | 34 |
| Pi09 | P07-B6 | 10 days | random | No | 813 | Frondihabitans peucedani RS-15(T) | 99 | 56 |
| Pi09 | P07-C1 | 10 days | random | No | 541 | Frondihabitans cladoniiphilus CafT13(T) | 98 | 38 |
| Pi09 | P07-C2 | 10 days | random | No | 905 | Frondihabitans peucedani RS-15(T) | 100 | 62 |
| Pi09 | P07-C3 | 10 days | random | No | 847 | Frondihabitans peucedani RS-15(T) | 100 | 58 |
| Pi09 | P07-C5 | 10 days | random | No | 901 | Frondihabitans peucedani RS-15(T) | 100 | 62 |
| Pi09 | P07-C6 | 10 days | random | No | 680 | Frondihabitans cladoniiphilus CafT13(T) | 99 | 47 |
| Pi09 | P07-D1 | 10 days | random | No | 713 | Frondihabitans cladoniiphilus CafT13(T) | 98 | 49 |
| Pi09 | P07-D2 | 10 days | random | No | 557 | Frondihabitans cladoniiphilus CafT13(T) | 98 | 38 |
| Pi09 | P07-D3 | 10 days | random | No | 800 | Frondihabitans cladoniiphilus CafT13(T) | 100 | 55 |
| Pi09 | P07-D4 | 10 days | random | No | 602 | Frondihabitans cladoniiphilus CafT13(T) | 100 | 42 |
| Pi09 | P07-D5 | 10 days | random | No | 870 | Frondihabitans peucedani RS-15(T) | 100 | 60 |
| Pi09 | P07-D6 | 10 days | random | No | 801 | Frondihabitans cladoniiphilus CafT13(T) | 100 | 55 |
| Pi09 | P41-B2 | 3 weeks | random | No | 1051 | Gryllotalpicola ginsengisoli DSM 22003(T) | 98 | 73 |
| Pi10 | P21-D4 | 10 days | random | No | 965 | Sphingomonas ginsenosidimutans Gsoil 1429(T) | 97 | 68 |
| Pi10 | P21-D5 | 10 days | random | No | 658 | Sphingomonas oligoaromativorans SY-6(T) | 99 | 47 |
| Pi10 | P21-D6 | 10 days | random | No | 1074 | Burkholderia udeis LMG 27134(T) | 99 | 74 |
| Pi11 | P08-A1 | 10 days | random | No | 1063 | Phenylobacterium lituiforme Fail3(T) | 96 | 76 |

| | | | | | | | | |
|---|---|---|---|---|---|---|---|---|
| Pi11 | P08-A2 | 10 days | random | No | 1080 | Phenylobacterium lituiforme Fail3(T) | 96 | 77 |
| Pi11 | P08-A3 | 10 days | random | No | 1071 | Phenylobacterium lituiforme Fail3(T) | 96 | 76 |
| Pi11 | P08-A4 | 10 days | random | No | 1139 | Phenylobacterium lituiforme Fail3(T) | 96 | 81 |
| Pi11 | P08-A5 | 10 days | random | No | 1153 | Phenylobacterium lituiforme Fail3(T) | 96 | 82 |
| Pi11 | P08-A6 | 10 days | random | No | 1084 | Phenylobacterium lituiforme Fail3(T) | 96 | 77 |
| Pi11 | P08-B1 | 10 days | random | No | 858 | Phenylobacterium lituiforme Fail3(T) | 96 | 61 |
| Pi11 | P08-B2 | 10 days | random | No | 1082 | Phenylobacterium lituiforme Fail3(T) | 96 | 77 |
| Pi11 | P08-B3 | 10 days | random | No | 1078 | Burkholderia sordidicola S5-B(T) | 98 | 74 |
| Pi11 | P08-B4 | 10 days | random | No | 1039 | Phenylobacterium lituiforme Fail3(T) | 96 | 74 |
| Pi11 | P08-B5 | 10 days | random | No | 1169 | Caulobacter mirabilis FWC38(T) | 96 | 83 |
| Pi11 | P08-B6 | 10 days | random | No | 994 | Phenylobacterium lituiforme Fail3(T) | 96 | 71 |
| Pi11 | P08-C1 | 10 days | random | No | 1079 | Phenylobacterium lituiforme Fail3(T) | 96 | 77 |
| Pi11 | P08-C2 | 10 days | random | No | 619 | Phenylobacterium lituiforme Fail3(T) | 96 | 45 |
| Pi11 | P08-C3 | 10 days | random | No | 1077 | Phenylobacterium lituiforme Fail3(T) | 96 | 77 |
| Pi11 | P08-C4 | 10 days | random | No | 1139 | Caulobacter mirabilis FWC38(T) | 96 | 81 |
| Pi11 | P08-C5 | 10 days | random | No | 910 | Caulobacter mirabilis FWC38(T) | 94 | 65 |
| Pi11 | P08-C6 | 10 days | random | No | 1057 | Phenylobacterium lituiforme Fail3(T) | 96 | 75 |
| Pi11 | P08-D1 | 10 days | random | No | 1148 | Phenylobacterium lituiforme Fail3(T) | 96 | 82 |
| Pi11 | P08-D2 | 10 days | random | No | 1159 | Phenylobacterium lituiforme Fail3(T) | 96 | 83 |
| Pi11 | P08-D3 | 10 days | random | No | 1018 | Phenylobacterium lituiforme Fail3(T) | 96 | 73 |
| Pi11 | P08-D4 | 10 days | random | No | 1098 | Phenylobacterium lituiforme Fail3(T) | 96 | 78 |
| Pi11 | P08-D5 | 10 days | random | No | 930 | Phenylobacterium lituiforme Fail3(T) | 97 | 66 |
| Pi11 | P08-D6 | 10 days | random | No | 1145 | Phenylobacterium lituiforme Fail3(T) | 96 | 82 |
| Pi11 | P28-A1 | 3 weeks | random | No | 625 | Bradyrhizobium iriomotense EK05(T) | 99 | 44 |
| Pi11 | P28-A2 | 3 weeks | random | No | 970 | Methylovirgula ligni BW863(T) | 99 | 69 |
| Pi11 | P28-A3 | 3 weeks | random | No | 924 | Methylovirgula ligni BW863(T) | 99 | 66 |
| Pi11 | P28-A4 | 3 weeks | random | No | 903 | Acidisphaera rubrifaciens HS-AP3(T) | 95 | 64 |
| Pi11 | P28-A5 | 3 weeks | random | No | 464 | Acidobacterium capsulatum ATCC 51196(T) | 94 | 33 |
| Pi11 | P28-A6 | 3 weeks | random | No | 630 | Acidobacterium capsulatum ATCC 51196(T) | 97 | 44 |
| Pi11 | P28-B1 | 3 weeks | random | No | 914 | Methylovirgula ligni BW863(T) | 99 | 65 |
| Pi11 | P28-B2 | 3 weeks | random | No | 1066 | Acidisoma tundrae WM1(T) | 99 | 75 |
| Pi11 | P28-B3 | 3 weeks | random | No | 1128 | Brevundimonas aveniformis EMB102(T) | 96 | 81 |
| Pi11 | P28-B4 | 3 weeks | random | No | 931 | Methylovirgula ligni BW863(T) | 99 | 66 |

| | | | | | | | | |
|---|---|---|---|---|---|---|---|---|
| Pi11 | P28-B5 | 3 weeks | random | No | 940 | Acidisphaera rubrifaciens HS-AP3(T) | 95 | 67 |
| Pi11 | P28-C1 | 3 weeks | random | No | 728 | Methylovirgula ligni BW863(T) | 98 | 52 |
| Pi11 | P28-C2 | 3 weeks | random | No | 891 | Phenylobacterium lituiforme FaiI3(T) | 96 | 64 |
| Pi11 | P28-C3 | 3 weeks | random | No | 958 | Methylovirgula ligni BW863(T) | 99 | 68 |
| Pi11 | P28-C4 | 3 weeks | random | No | 964 | Phenylobacterium lituiforme FaiI3(T) | 92 | 68 |
| Pi11 | P28-C5 | 3 weeks | random | No | 936 | Methylovirgula ligni BW863(T) | 99 | 67 |
| Pi11 | P28-C6 | 3 weeks | random | No | 1023 | Acidisphaera rubrifaciens HS-AP3(T) | 95 | 72 |
| Pi11 | P28-D1 | 3 weeks | random | No | 926 | Methylovirgula ligni BW863(T) | 99 | 66 |
| Pi11 | P28-D3 | 3 weeks | random | No | 926 | Acidisphaera rubrifaciens HS-AP3(T) | 95 | 66 |
| Pi11 | P28-D6 | 3 weeks | random | No | 996 | Granulicella rosea TPO1014(T) | 98 | 70 |
| Pi11 | P46-A1 | 6 weeks | random | No | 476 | Methyloferula stellata AR4(T) | 99 | 34 |
| Pi11 | P46-A2 | 6 weeks | random | No | 771 | Methyloferula stellata AR4(T) | 98 | 55 |
| Pi11 | P46-A3 | 6 weeks | random | No | 620 | Gluconacetobacter takamatsuzukensis T61213-20-1a(T) | 97 | 44 |
| Pi11 | P46-A4 | 6 weeks | random | No | 882 | 'Acidipila rosea' AP8(T) | 97 | 63 |
| Pi11 | P46-A6 | 6 weeks | random | No | 299 | Methyloferula stellata AR4(T) | 99 | 21 |
| Pi11 | P46-B1 | 6 weeks | random | No | 762 | Methyloferula stellata AR4(T) | 98 | 54 |
| Pi11 | P46-B2 | 6 weeks | random | No | 769 | Methyloferula stellata AR4(T) | 98 | 55 |
| Pi11 | P46-B3 | 6 weeks | random | No | 944 | Granulicella arctica MP5ACTX2(T) | 99 | 66 |
| Pi11 | P46-B4 | 6 weeks | random | No | 724 | 'Acidipila rosea' AP8(T) | 96 | 51 |
| Pi12 | P01-A1 | 10 days | random | No | 867 | Burkholderia sordidicola S5-B(T) | 100 | 60 |
| Pi12 | P01-A2 | 10 days | random | No | 892 | Burkholderia udeis LMG 27134(T) | 99 | 61 |
| Pi12 | P01-A3 | 10 days | random | No | 823 | Granulicella pectinivorans TPB6011(T) | 98 | 58 |
| Pi12 | P01-A4 | 10 days | random | No | 249 | Burkholderia udeis LMG 27134(T) | 100 | 17 |
| Pi12 | P01-A5 | 10 days | random | No | 172 | Acidisoma tundrae WM1(T) | 97 | 12 |
| Pi12 | P01-A6 | 10 days | random | No | 867 | Burkholderia phenazinium LMG 2247(T) | 99 | 60 |
| Pi12 | P01-B1 | 10 days | random | No | 894 | Sphingomonas oligoaromativorans SY-6(T) | 98 | 63 |
| Pi12 | P01-B2 | 10 days | random | No | 864 | Burkholderia phenazinium LMG 2247(T) | 99 | 59 |
| Pi12 | P01-B3 | 10 days | random | No | 831 | Burkholderia sordidicola S5-B(T) | 99 | 57 |
| Pi12 | P01-B4 | 10 days | random | No | 842 | Burkholderia sordidicola S5-B(T) | 99 | 58 |
| Pi12 | P01-B5 | 10 days | random | No | 922 | Phenylobacterium lituiforme FaiI3(T) | 97 | 66 |
| Pi12 | P01-B6 | 10 days | random | No | 802 | Burkholderia phenazinium LMG 2247(T) | 99 | 55 |
| Pi12 | P01-C1 | 10 days | random | No | 548 | Gluconacetobacter asukensis K8617-1-1b(T) | 94 | 39 |
| Pi12 | P01-C2 | 10 days | random | No | 936 | 'Acidipila rosea' AP8(T) | 98 | 66 |

| | | | | | | | | |
|---|---|---|---|---|---|---|---|---|
| Pi12 | P01-C3 | 10 days | random | No | 807 | Burkholderia sordidicola S5-B(T) | 99 | 56 |
| Pi12 | P01-C4 | 10 days | random | No | 871 | Burkholderia sordidicola S5-B(T) | 99 | 60 |
| Pi12 | P01-C5 | 10 days | random | No | 899 | Burkholderia udeis LMG 27134(T) | 99 | 62 |
| Pi12 | P01-C6 | 10 days | random | No | 879 | Burkholderia udeis LMG 27134(T) | 99 | 61 |
| Pi12 | P01-D1 | 10 days | random | No | 821 | Granulicella pectinivorans TPB6011(T) | 98 | 58 |
| Pi12 | P01-D2 | 10 days | random | No | 834 | Granulicella pectinivorans TPB6011(T) | 98 | 59 |
| Pi12 | P01-D3 | 10 days | random | No | 867 | Burkholderia sordidicola S5-B(T) | 99 | 60 |
| Pi12 | P01-D4 | 10 days | random | No | 868 | Sphingomonas oligoaromativorans SY-6(T) | 98 | 62 |
| Pi12 | P01-D5 | 10 days | random | No | 871 | 'Acidipila rosea' AP8(T) | 98 | 61 |
| Pi12 | P01-D6 | 10 days | random | No | 679 | Burkholderia sordidicola S5-B(T) | 100 | 47 |
| Pi12 | P23-A2 | 3 weeks | random | No | 898 | Acidisphaera rubrifaciens HS-AP3(T) | 95 | 64 |
| Pi12 | P23-A3 | 3 weeks | random | No | 556 | Methylovirgula ligni BW863(T) | 98 | 40 |
| Pi12 | P23-A4 | 3 weeks | random | No | 272 | PAC000434_s | 99 | 19 |
| Pi12 | P23-A5 | 3 weeks | random | No | 645 | Granulicella arctica MP5ACTX2(T) | 99 | 47 |
| Pi12 | P23-A6 | 3 weeks | random | No | 693 | Acidisoma tundrae WM1(T) | 96 | 49 |
| Pi12 | P23-B1 | 3 weeks | random | No | 509 | Tanticharoenia sakaeratensis NBRC 103193(T) | 97 | 37 |
| Pi12 | P23-B2 | 3 weeks | random | No | 785 | Methylovirgula ligni BW863(T) | 98 | 56 |
| Pi12 | P23-B3 | 3 weeks | random | No | 927 | Acidisphaera rubrifaciens HS-AP3(T) | 95 | 66 |
| Pi12 | P23-B4 | 3 weeks | random | No | 833 | Acidisphaera rubrifaciens HS-AP3(T) | 96 | 59 |
| Pi12 | P23-B5 | 3 weeks | random | No | 823 | Acidisphaera rubrifaciens HS-AP3(T) | 96 | 58 |
| Pi12 | P23-B6 | 3 weeks | random | No | 968 | Acidisphaera rubrifaciens HS-AP3(T) | 95 | 68 |
| Pi12 | P23-C1 | 3 weeks | random | No | 908 | Methylovirgula ligni BW863(T) | 99 | 65 |
| Pi12 | P23-C2 | 3 weeks | random | No | 971 | Acidisphaera rubrifaciens HS-AP3(T) | 96 | 69 |
| Pi12 | P23-C3 | 3 weeks | random | No | 920 | Beijerinckia derxii supsp. derxii DSM 2328(T) | 99 | 66 |
| Pi12 | P23-C4 | 3 weeks | random | No | 964 | Acidisphaera rubrifaciens HS-AP3(T) | 95 | 68 |
| Pi12 | P23-C5 | 3 weeks | random | No | 416 | Labedella gwakjiensis KSW2-17(T) | 99 | 29 |
| Pi12 | P23-C6 | 3 weeks | random | No | 962 | Acidisphaera rubrifaciens HS-AP3(T) | 95 | 68 |
| Pi12 | P23-D1 | 3 weeks | random | No | 794 | Methylovirgula ligni BW863(T) | 99 | 57 |
| Pi12 | P23-D2 | 3 weeks | random | No | 832 | Methylovirgula ligni BW863(T) | 98 | 59 |
| Pi12 | P23-D3 | 3 weeks | random | No | 828 | Acidisphaera rubrifaciens HS-AP3(T) | 96 | 58 |
| Pi12 | P23-D4 | 3 weeks | random | No | 950 | Methylovirgula ligni BW863(T) | 99 | 68 |
| Pi12 | P23-D5 | 3 weeks | random | No | 490 | Tanticharoenia sakaeratensis NBRC 103193(T) | 97 | 35 |
| Pi12 | P23-D6 | 3 weeks | random | No | 639 | Acidisoma tundrae WM1(T) | 99 | 45 |

| | | | | | | | | |
|---|---|---|---|---|---|---|---|---|
| Pi13 | P10-A1 | 10 days | morphology | No | 646 | Burkholderia phenazinium LMG 2247(T) | 98 | 45 |
| Pi13 | P10-A2 | 10 days | random | No | 422 | Rhodanobacter ginsenosidimutans Gsoil 3054(T) | 99 | 29 |
| Pi13 | P10-A3 | 10 days | random | No | 272 | Mycobacterium rhodesiae DSM 44223(T) | 100 | 19 |
| Pi13 | P10-A4 | 10 days | random | No | 1132 | Phenylobacterium composti 4T-6(T) | 96 | 81 |
| Pi13 | P10-A6 | 10 days | random | No | 1023 | Rhodanobacter umsongensis GR24-2(T) | 99 | 70 |
| Pi13 | P10-B1 | 10 days | random | No | 1022 | Rhodanobacter umsongensis GR24-2(T) | 99 | 70 |
| Pi13 | P10-B2 | 10 days | random | No | 1050 | Dyella japonica XD53(T) | 100 | 72 |
| Pi13 | P10-B3 | 10 days | random | No | 1104 | Acidisoma tundrae WM1(T) | 99 | 78 |
| Pi13 | P10-B4 | 10 days | random | No | 1103 | Burkholderia sordidicola S5-B(T) | 98 | 76 |
| Pi13 | P10-B5 | 10 days | random | No | 925 | Microbacterium immunditiarum SK 18(T) | 99 | 64 |
| Pi13 | P10-B6 | 10 days | random | No | 999 | Dyella japonica XD53(T) | 100 | 68 |
| Pi13 | P10-C1 | 10 days | random | No | 1024 | Rhodanobacter umsongensis GR24-2(T) | 99 | 70 |
| Pi13 | P10-C2 | 10 days | random | No | 1076 | Phenylobacterium composti 4T-6(T) | 96 | 77 |
| Pi13 | P10-C3 | 10 days | random | No | 1064 | Phenylobacterium composti 4T-6(T) | 96 | 76 |
| Pi13 | P10-C4 | 10 days | random | No | 1024 | Rhodanobacter umsongensis GR24-2(T) | 99 | 70 |
| Pi13 | P10-C5 | 10 days | random | No | 1024 | Dyella koreensis BB4(T) | 99 | 70 |
| Pi13 | P10-C6 | 10 days | random | No | 1046 | Dyella japonica XD53(T) | 100 | 71 |
| Pi13 | P10-D1 | 10 days | random | No | 1108 | Rhodanobacter umsongensis GR24-2(T) | 99 | 76 |
| Pi13 | P10-D2 | 10 days | random | No | 1127 | Phenylobacterium composti 4T-6(T) | 96 | 80 |
| Pi13 | P10-D3 | 10 days | random | No | 484 | Acidisoma sibiricum TPB606(T) | 97 | 34 |
| Pi13 | P10-D4 | 10 days | random | No | 830 | Mycobacterium neoaurum ATCC 25795(T) | 99 | 58 |
| Pi13 | P10-D5 | 10 days | random | No | 1141 | Phenylobacterium lituiforme FaiI3(T) | 95 | 81 |
| Pi13 | P10-D6 | 10 days | random | No | 1024 | Dyella ginsengisoli Gsoil 3046(T) | 99 | 70 |
| Pi13 | P30-A1 | 3 weeks | random | No | 1029 | Acidisoma sibiricum TPB606(T) | 99 | 73 |
| Pi13 | P30-A2 | 3 weeks | random | No | 1039 | Marmoricola aequoreus SST-45(T) | 98 | 72 |
| Pi13 | P30-A4 | 3 weeks | random | No | 598 | Mycobacterium abscessus subsp. bolletii BD(T) | 100 | 42 |
| Pi13 | P30-A5 | 3 weeks | random | No | 581 | Rhodanobacter ginsengisoli GR17-7(T) | 99 | 40 |
| Pi13 | P30-A6 | 3 weeks | random | No | 848 | Diaminobutyricimonas aerilata 6408J-67(T) | 98 | 59 |
| Pi13 | P30-B1 | 3 weeks | random | No | 608 | Mycobacterium austroafricanum ATCC 33464(T) | 98 | 42 |
| Pi13 | P30-B2 | 3 weeks | random | No | 1051 | Dyella japonica XD53(T) | 100 | 72 |
| Pi13 | P30-B3 | 3 weeks | random | No | 895 | Rhodanobacter ginsengisoli GR17-7(T) | 99 | 61 |
| Pi13 | P30-B4 | 3 weeks | random | No | 914 | Granulicella arctica MP5ACTX2(T) | 97 | 64 |
| Pi13 | P30-B5 | 3 weeks | random | No | 911 | Rhodanobacter ginsengisoli GR17-7(T) | 99 | 62 |

| | | | | | | | | |
|---|---|---|---|---|---|---|---|---|
| Pi13 | P30-B6 | 3 weeks | random | No | 1044 | Gluconacetobacter asukensis K8617-1-1b(T) | 95 | 74 |
| Pi13 | P30-C1 | 3 weeks | random | No | 645 | Methylovirgula ligni BW863(T) | 98 | 46 |
| Pi13 | P30-C2 | 3 weeks | random | No | 858 | Methylovirgula ligni BW863(T) | 98 | 61 |
| Pi13 | P30-C3 | 3 weeks | random | No | 991 | Rhodanobacter ginsengisoli GR17-7(T) | 98 | 68 |
| Pi13 | P30-C4 | 3 weeks | random | No | 713 | Methylovirgula ligni BW863(T) | 97 | 51 |
| Pi13 | P30-C5 | 3 weeks | random | No | 800 | Methylocella tundrae T4(T) | 97 | 57 |
| Pi13 | P30-C6 | 3 weeks | random | No | 688 | Methylovirgula ligni BW863(T) | 99 | 49 |
| Pi13 | P30-D1 | 3 weeks | random | No | 1159 | Acidisphaera rubrifaciens HS-AP3(T) | 96 | 82 |
| Pi13 | P30-D2 | 3 weeks | random | No | 612 | Methylovirgula ligni BW863(T) | 98 | 44 |
| Pi13 | P30-D4 | 3 weeks | random | No | 891 | Rhodanobacter ginsengisoli GR17-7(T) | 99 | 61 |
| Pi13 | P30-D5 | 3 weeks | random | No | 826 | Rhodanobacter ginsengisoli GR17-7(T) | 99 | 56 |
| Pi13 | P30-D6 | 3 weeks | random | No | 645 | Mycobacterium austroafricanum ATCC 33464(T) | 98 | 45 |
| Pi13 | P45-C2 | 6 weeks | random | No | 913 | Methylovirgula ligni BW863(T) | 99 | 65 |
| Pi13 | P45-C3 | 6 weeks | random | No | 326 | Methylocapsa aurea KYG(T) | 97 | 23 |
| Pi13 | P45-C4 | 6 weeks | random | No | 811 | Bradyrhizobium paxllaeri LMTR 21(T) | 99 | 58 |
| Pi13 | P45-C5 | 6 weeks | random | No | 656 | Methylovirgula ligni BW863(T) | 98 | 47 |
| Pi13 | P45-D4 | 6 weeks | random | No | 744 | Labedella gwakjiensis KSW2-17(T) | 99 | 51 |
| Pi13 | P45-D5 | 6 weeks | random | No | 894 | Mycobacterium abscessus subsp. bolletii BD(T) | 100 | 62 |
| Pi13 | P45-D6 | 6 weeks | random | No | 1007 | Mycobacterium abscessus subsp, bolletii BD(T) | 100 | 70 |
| Pi14 | P06-A1 | 10 days | random | No | 922 | Burkholderia udeis LMG 27134(T) | 99 | 63 |
| Pi14 | P06-A2 | 10 days | random | No | 789 | Cryobacterium mesophilum MSL-15(T) | 99 | 55 |
| Pi14 | P06-A3 | 10 days | random | No | 917 | Burkholderia phenazinium LMG 2247(T) | 100 | 63 |
| Pi14 | P06-A4 | 10 days | random | No | 560 | Streptacidiphilus albus DSM 41753(T) | 97 | 40 |
| Pi14 | P06-A5 | 10 days | random | No | 479 | Burkholderia phenazinium LMG 2247(T) | 99 | 33 |
| Pi14 | P06-A6 | 10 days | random | No | 774 | Burkholderia sordidicola S5-B(T) | 98 | 53 |
| Pi14 | P06-B3 | 10 days | random | No | 314 | Burkholderia phenazinium LMG 2247(T) | 99 | 22 |
| Pi14 | P06-B4 | 10 days | random | No | 741 | Burkholderia phenazinium LMG 2247(T) | 100 | 51 |
| Pi14 | P06-B5 | 10 days | random | No | 818 | Burkholderia sordidicola S5-B(T) | 99 | 56 |
| Pi14 | P06-B6 | 10 days | random | No | 714 | Burkholderia phenazinium LMG 2247(T) | 99 | 49 |
| Pi14 | P06-C1 | 10 days | random | No | 466 | Dyella marensis CS5-B2(T) | 98 | 32 |
| Pi14 | P06-C2 | 10 days | random | No | 806 | Luteimicrobium album RI148-Li105(T) | 100 | 56 |
| Pi14 | P06-C3 | 10 days | random | No | 421 | Burkholderia udeis LMG 27134(T) | 100 | 29 |
| Pi14 | P06-C4 | 10 days | random | No | 868 | Burkholderia sordidicola S5-B(T) | 99 | 60 |

| | | | | | | | | |
|---|---|---|---|---|---|---|---|---|
| Pi14 | P06-C5 | 10 days | random | No | 818 | Burkholderia sordidicola S5-B(T) | 99 | 56 |
| Pi14 | P06-C6 | 10 days | random | No | 817 | Burkholderia phenazinium LMG 2247(T) | 99 | 56 |
| Pi14 | P06-D1 | 10 days | random | No | 738 | Burkholderia sordidicola S5-B(T) | 99 | 51 |
| Pi14 | P06-D2 | 10 days | random | No | 877 | Rhodanobacter umsongensis GR24-2(T) | 99 | 60 |
| Pi14 | P06-D3 | 10 days | random | No | 503 | Burkholderia udeis LMG 27134(T) | 100 | 35 |
| Pi14 | P06-D4 | 10 days | random | No | 647 | Burkholderia udeis LMG 27134(T) | 99 | 45 |
| Pi14 | P06-D5 | 10 days | random | No | 740 | Burkholderia sordidicola S5-B(T) | 99 | 51 |
| Pi14 | P06-D6 | 10 days | random | No | 724 | Burkholderia udeis LMG 27134(T) | 98 | 50 |
| Pi14 | P27-A4 | 3 weeks | random | No | 800 | Bradyrhizobium iriomotense EK05(T) | 98 | 57 |
| Pi14 | P27-A5 | 3 weeks | random | No | 1007 | Methylovirgula ligni BW863(T) | 99 | 72 |
| Pi14 | P27-B1 | 3 weeks | random | No | 640 | Rhodanobacter umsongensis GR24-2(T) | 99 | 44 |
| Pi14 | P27-B3 | 3 weeks | random | No | 1023 | Gluconacetobacter sacchari SRI 1794(T) | 96 | 72 |
| Pi14 | P27-B5 | 3 weeks | random | No | 511 | Rhodanobacter umsongensis GR24-2(T) | 99 | 35 |
| Pi14 | P27-B6 | 3 weeks | random | No | 749 | Bradyrhizobium iriomotense EK05(T) | 99 | 53 |
| Pi14 | P27-C1 | 3 weeks | random | No | 984 | Cryobacterium mesophilum MSL-15(T) | 98 | 68 |
| Pi14 | P27-C2 | 3 weeks | random | No | 1076 | Acidisphaera rubrifaciens HS-AP3(T) | 96 | 76 |
| Pi14 | P27-C5 | 3 weeks | random | No | 749 | Bradyrhizobium paxllaeri LMTR 21(T) | 100 | 53 |
| Pi14 | P27-C6 | 3 weeks | random | No | 1000 | Rhodanobacter umsongensis GR24-2(T) | 99 | 68 |
| Pi14 | P27-D1 | 3 weeks | random | No | 664 | Rhodanobacter ginsengisoli GR17-7(T) | 100 | 45 |
| Pi14 | P27-D2 | 3 weeks | random | No | 623 | Dyella kyungheensis THG-B117(T) | 99 | 42 |
| Pi14 | P27-D4 | 3 weeks | random | No | 645 | Methylovirgula ligni BW863(T) | 98 | 46 |
| Pi14 | P27-D5 | 3 weeks | random | No | 889 | Gryllotalpicola daejeonensis RU-04(T) | 99 | 61 |
| Pi14 | P44-C1 | 6 weeks | random | No | 885 | Methylovirgula ligni BW863(T) | 99 | 63 |
| Pi14 | P44-C2 | 6 weeks | random | No | 794 | Labedella gwakjiensis KSW2-17(T) | 99 | 55 |
| Pi14 | P44-C3 | 6 weeks | random | No | 968 | Methylovirgula ligni BW863(T) | 99 | 69 |
| Pi14 | P44-C4 | 6 weeks | random | No | 881 | Kozakia baliensis Yo-3(T) | 96 | 62 |
| Pi14 | P44-C5 | 6 weeks | random | No | 575 | Methylovirgula ligni BW863(T) | 98 | 41 |
| Pi14 | P44-C6 | 6 weeks | random | No | 816 | Acidisoma tundrae WM1(T) | 99 | 58 |
| Pi14 | P44-D1 | 6 weeks | random | No | 505 | Acidisoma sibiricum TPB606(T) | 95 | 36 |
| Pi14 | P44-D2 | 6 weeks | random | No | 979 | Gluconacetobacter liquefaciens IFO 12388(T) | 95 | 68 |
| Pi14 | P44-D4 | 6 weeks | random | No | 776 | Kozakia baliensis Yo-3(T) | 97 | 55 |
| Pi14 | P44-D6 | 6 weeks | random | No | 1014 | Bradyrhizobium elkanii USDA 76(T) | 99 | 72 |
| Pi16 | P05-A2 | 10 days | morphology | No | 736 | Burkholderia sordidicola S5-B(T) | 99 | 51 |

| | | | | | | | | |
|---|---|---|---|---|---|---|---|---|
| Pi16 | P05-A3 | 10 days | morphology | No | 395 | Sphingomonas oligoaromativorans SY-6(T) | 99 | 28 |
| Pi16 | P05-A5 | 10 days | morphology | No | 54 | Burkholderia zhejiangensis OP-1(T) | 98 | 4 |
| Pi16 | P05-A6 | 10 days | morphology | No | 722 | 'Acidipila rosea' AP8(T) | 97 | 51 |
| Pi16 | P05-B2 | 10 days | morphology | No | 499 | Bradyrhizobium elkanii USDA 76(T) | 98 | 35 |
| Pi16 | P05-B3 | 10 days | random | No | 433 | Burkholderia udeis LMG 27134(T) | 100 | 30 |
| Pi16 | P05-B6 | 10 days | random | No | 766 | Burkholderia sordidicola S5-B(T) | 99 | 53 |
| Pi16 | P05-C1 | 10 days | random | No | 699 | Burkholderia udeis LMG 27134(T) | 98 | 48 |
| Pi16 | P05-C2 | 10 days | random | No | 459 | Burkholderia phenazinium LMG 2247(T) | 99 | 32 |
| Pi16 | P05-C3 | 10 days | random | No | 242 | Burkholderia sordidicola S5-B(T) | 100 | 17 |
| Pi16 | P05-C4 | 10 days | random | No | 376 | Burkholderia sordidicola S5-B(T) | 100 | 26 |
| Pi16 | P05-C5 | 10 days | random | No | 598 | Burkholderia sordidicola S5-B(T) | 99 | 41 |
| Pi16 | P05-C6 | 10 days | random | No | 696 | Burkholderia udeis LMG 27134(T) | 99 | 48 |
| Pi16 | P05-D1 | 10 days | random | No | 735 | Burkholderia udeis LMG 27134(T) | 99 | 51 |
| Pi16 | P05-D2 | 10 days | random | No | 692 | Burkholderia udeis LMG 27134(T) | 99 | 48 |
| Pi16 | P05-D4 | 10 days | random | No | 442 | Burkholderia udeis LMG 27134(T) | 100 | 31 |
| Pi16 | P05-D5 | 10 days | random | No | 498 | Burkholderia udeis LMG 27134(T) | 99 | 34 |
| Pi16 | P05-D6 | 10 days | random | No | 760 | Burkholderia udeis LMG 27134(T) | 99 | 52 |
| Pi16 | P26-A1 | 3 weeks | random | No | 565 | Curtobacterium luteum DSM 20542(T) | 98 | 39 |
| Pi16 | P26-A2 | 3 weeks | random | No | 1042 | Acidisoma tundrae WM1(T) | 99 | 73 |
| Pi16 | P26-A3 | 3 weeks | random | No | 1024 | Acidisphaera rubrifaciens HS-AP3(T) | 95 | 73 |
| Pi16 | P26-A4 | 3 weeks | random | No | 1042 | Acidisoma tundrae WM1(T) | 99 | 73 |
| Pi16 | P26-A5 | 3 weeks | random | No | 1047 | Granulicella pectinivorans TPB6011(T) | 97 | 74 |
| Pi16 | P26-B1 | 3 weeks | random | No | 702 | Granulicella arctica MP5ACTX2(T) | 99 | 49 |
| Pi16 | P26-B2 | 3 weeks | random | No | 1069 | Acidisoma tundrae WM1(T) | 99 | 75 |
| Pi16 | P26-B3 | 3 weeks | random | No | 1135 | Granulibacter bethesdensis CGDNIH1(T) | 96 | 80 |
| Pi16 | P26-B4 | 3 weeks | random | Dyella, Burkholderia | 1133 | Acidisoma tundrae WM1(T) | 99 | 80 |
| Pi16 | P26-B5 | 3 weeks | random | No | 1100 | 'Acidipila rosea' AP8(T) | 97 | 77 |
| Pi16 | P26-B6 | 3 weeks | random | Dyella, Burkholderia | 598 | Acidisoma tundrae WM1(T) | 98 | 42 |
| Pi16 | P26-C1 | 3 weeks | random | No | 1025 | Granulicella arctica MP5ACTX2(T) | 98 | 72 |
| Pi16 | P26-C2 | 3 weeks | random | Dyella, Burkholderia | 1067 | Acidisoma tundrae WM1(T) | 99 | 75 |
| Pi16 | P26-C3 | 3 weeks | random | No | 861 | Granulicella aggregans TPB6028(T) | 98 | 62 |
| Pi16 | P26-C4 | 3 weeks | random | No | 817 | Acidisoma tundrae WM1(T) | 95 | 58 |
| Pi16 | P26-C5 | 3 weeks | random | Dyella | 1012 | Flavobacterium oceanosedimentum ATCC 31317(T) | 97 | 70 |

| | | | | | | | | |
|---|---|---|---|---|---|---|---|---|
| Pi16 | P26-C6 | 3 weeks | random | No | 1145 | Granulibacter bethesdensis CGDNIH1(T) | 96 | 81 |
| Pi16 | P26-D2 | 3 weeks | random | No | 1060 | 'Acidipila rosea' AP8(T) | 98 | 75 |
| Pi16 | P26-D3 | 3 weeks | random | No | 1029 | Acidisoma tundrae WM1(T) | 99 | 73 |
| Pi16 | P26-D4 | 3 weeks | random | No | 1092 | Acidisoma tundrae WM1(T) | 99 | 77 |
| Pi16 | P26-D5 | 3 weeks | random | No | 976 | Acidisphaera rubrifaciens HS-AP3(T) | 96 | 69 |
| Pi16 | P26-D6 | 3 weeks | random | No | 937 | Acidisphaera rubrifaciens HS-AP3(T) | 95 | 66 |
| Pi16 | P44-A1 | 6 weeks | random | No | 948 | Acidicapsa ligni WH120(T) | 99 | 67 |
| Pi16 | P44-A3 | 6 weeks | random | No | 773 | Acidisoma tundrae WM1(T) | 97 | 55 |
| Pi16 | P44-A4 | 6 weeks | random | No | 737 | Acidisoma tundrae WM1(T) | 96 | 52 |
| Pi16 | P44-A6 | 6 weeks | random | No | 542 | Granulicella pectinivorans TPB6011(T) | 99 | 38 |
| Pi16 | P44-B1 | 6 weeks | random | No | 930 | Granulicella aggregans TPB6028(T) | 98 | 65 |
| Pi16 | P44-B2 | 6 weeks | random | No | 908 | Acidisphaera rubrifaciens HS-AP3(T) | 95 | 64 |
| Pi16 | P44-B5 | 6 weeks | random | No | 568 | Methylovirgula ligni BW863(T) | 98 | 41 |
| Pi16 | P44-B6 | 6 weeks | random | No | 856 | Methylovirgula ligni BW863(T) | 98 | 61 |
| Pi17 | P03-A1 | 10 days | morphology | No | 1025 | Burkholderia xenovorans LB400(T) | 99 | 71 |
| Pi17 | P03-A2 | 10 days | morphology | No | 819 | Frondihabitans peucedani RS-15(T) | 100 | 56 |
| Pi17 | P03-A3 | 10 days | morphology | No | 812 | Dyella japonica XD53(T) | 99 | 55 |
| Pi17 | P03-A4 | 10 days | morphology | No | 669 | Dyella marensis CS5-B2(T) | 99 | 46 |
| Pi17 | P03-A5 | 10 days | morphology | Staphylococcus, Dyella | 869 | Streptacidiphilus anmyonensis AM-11(T) | 99 | 60 |
| Pi17 | P03-A6 | 10 days | morphology | No | 413 | Burkholderia tuberum STM678(T) | 97 | 28 |
| Pi17 | P03-B1 | 10 days | random | No | 246 | Rhodanobacter panaciterrae LnR5-47(T) | 99 | 17 |
| Pi17 | P03-B2 | 10 days | random | No | 877 | Rhodanobacter umsongensis GR24-2(T) | 98 | 60 |
| Pi17 | P03-B3 | 10 days | random | No | 805 | Burkholderia bryophila LMG 23644(T) | 99 | 55 |
| Pi17 | P03-B4 | 10 days | random | No | 670 | Burkholderia sabiae Br3407(T) | 98 | 46 |
| Pi17 | P03-B5 | 10 days | random | No | 831 | Rhodanobacter umsongensis GR24-2(T) | 99 | 57 |
| Pi17 | P03-B6 | 10 days | random | No | 807 | Burkholderia graminis C4D1M(T) | 99 | 56 |
| Pi17 | P03-C1 | 10 days | random | No | 828 | Dyella soli JS12-10(T) | 98 | 56 |
| Pi17 | P03-C2 | 10 days | random | No | 664 | Burkholderia sordidicola S5-B(T) | 98 | 46 |
| Pi17 | P03-C3 | 10 days | random | No | 635 | Frateuria terrea VA24(T) | 97 | 46 |
| Pi17 | P03-C4 | 10 days | random | No | 694 | Mycobacterium fluoranthenivorans FA-4(T) | 99 | 48 |
| Pi17 | P03-C5 | 10 days | random | No | 1012 | Burkholderia phenazinium LMG 2247(T) | 95 | 70 |
| Pi17 | P03-C6 | 10 days | random | No | 958 | Burkholderia phenazinium LMG 2247(T) | 97 | 66 |
| Pi17 | P03-D1 | 10 days | random | No | 707 | Rhodanobacter umsongensis GR24-2(T) | 99 | 48 |

| | | | | | | | | |
|---|---|---|---|---|---|---|---|---|
| Pi17 | P43-D1 | 6 weeks | random | No | 697 | Methylovirgula ligni BW863(T) | 99 | 50 |
| Pi17 | P43-D2 | 6 weeks | random | No | 803 | Methylovirgula ligni BW863(T) | 99 | 57 |
| Pi17 | P43-D3 | 6 weeks | random | No | 800 | Pseudonocardia spinosispora LM 141(T) | 98 | 55 |
| Pi17 | P43-D4 | 6 weeks | random | Staphylococcus, Dyella | 511 | Streptacidiphilus anmyonensis AM-11(T) | 98 | 35 |
| Pi17 | P43-D5 | 6 weeks | random | No | 869 | 'Dyella jejuensis' JP1(T) | 99 | 59 |
| Pi17 | P43-D6 | 6 weeks | random | No | 759 | Labedella gwakjiensis KSW2-17(T) | 99 | 53 |
| Pi18 | P11-A1 | 10 days | morphology | No | 893 | Staphylococcus warneri ATCC 27836(T) | 99 | 61 |
| Pi18 | P11-A3 | 10 days | morphology | No | 1037 | Rhodanobacter fulvus Jip2(T) | 99 | 71 |
| Pi18 | P11-A5 | 10 days | morphology | No | 677 | Arthrobacter equi IMMIB L-1606(T) | 99 | 47 |
| Pi18 | P11-A6 | 10 days | morphology | No | 976 | Kitasatospora mediocidica IFO 14789(T) | 100 | 67 |
| Pi18 | P11-B1 | 10 days | morphology | No | 636 | Streptacidiphilus albus DSM 41753(T) | 99 | 44 |
| Pi18 | P11-B2 | 10 days | morphology | No | 1026 | Burkholderia bryophila LMG 23644(T) | 98 | 71 |
| Pi18 | P11-B3 | 10 days | random | No | 684 | Dyella soli JS12-10(T) | 99 | 47 |
| Pi18 | P11-C2 | 10 days | random | No | 671 | Dyella marensis CS5-B2(T) | 99 | 46 |
| Pi18 | P11-C4 | 10 days | random | No | 677 | Mycobacterium fluoranthenivorans FA-4(T) | 99 | 47 |
| Pi18 | P11-C5 | 10 days | random | No | 433 | Streptacidiphilus albus DSM 41753(T) | 99 | 30 |
| Pi18 | P11-C6 | 10 days | random | No | 1031 | Dyella ginsengisoli Gsoil 3046(T) | 99 | 70 |
| Pi18 | P11-D1 | 10 days | random | No | 782 | Mycobacterium fluoranthenivorans FA-4(T) | 99 | 54 |
| Pi18 | P11-D2 | 10 days | random | No | 692 | Mycobacterium mucogenicum ATCC 49650(T) | 99 | 48 |
| Pi18 | P11-D4 | 10 days | random | No | 891 | Burkholderia graminis C4D1M(T) | 99 | 61 |
| Pi18 | P11-D6 | 10 days | random | No | 819 | Mycobacterium phocaicum CIP 108542(T) | 99 | 57 |
| Pi18 | P31-A3 | 3 weeks | random | No | 308 | Actinospica robiniae GE134769(T) | 98 | 21 |
| Pi18 | P31-A4 | 3 weeks | random | No | 1000 | Rhodanobacter ginsengisoli GR17-7(T) | 99 | 68 |
| Pi18 | P31-A5 | 3 weeks | random | No | 650 | Rhodanobacter ginsengisoli GR17-7(T) | 100 | 44 |
| Pi18 | P31-B3 | 3 weeks | random | No | 616 | Rhodanobacter ginsengisoli GR17-7(T) | 100 | 42 |
| Pi18 | P31-C3 | 3 weeks | random | No | 877 | Mycobacterium mucogenicum ATCC 49650(T) | 99 | 61 |
| Pi18 | P31-D2 | 3 weeks | random | No | 782 | Streptacidiphilus durhamensis FSCA67(T) | 100 | 54 |
| Pi18 | P31-D4 | 3 weeks | random | No | 677 | Streptacidiphilus durhamensis FSCA67(T) | 100 | 47 |
| Pi18 | P46-C4 | 6 weeks | random | No | 440 | Methylovirgula ligni BW863(T) | 98 | 31 |
| Pi18 | P46-C5 | 6 weeks | random | No | 742 | Mycobacterium abscessus subsp, abscessus ATCC 1997 | 100 | 52 |
| Pi18 | P46-C6 | 6 weeks | random | No | 787 | Gluconacetobacter takamatsuzukensis T61213-20-1a(T | 96 | 56 |
| Pi19 | P02-A1 | 10 days | morphology | No | 840 | Rhodanobacter umsongensis GR24-2(T) | 99 | 57 |
| Pi19 | P02-A2 | 10 days | morphology | No | 863 | Dyella marensis CS5-B2(T) | 99 | 59 |

| | | | | | | | | |
|---|---|---|---|---|---|---|---|---|
| Pi19 | P02-A3 | 10 days | morphology | Dyella | 872 | Streptacidiphilus durhamensis FSCA67(T) | 100 | 61 |
| Pi19 | P02-A4 | 10 days | morphology | No | 768 | Streptacidiphilus durhamensis FSCA67(T) | 100 | 53 |
| Pi19 | P02-A5 | 10 days | morphology | No | 802 | Burkholderia sordidicola S5-B(T) | 99 | 55 |
| Pi19 | P02-A6 | 10 days | morphology | No | 834 | Dyella japonica XD53(T) | 100 | 57 |
| Pi19 | P02-B1 | 10 days | morphology | No | 774 | Leptothrix cholodnii CCM 1827 | 99 | 53 |
| Pi19 | P02-B2 | 10 days | morphology | No | 831 | Rhodanobacter umsongensis GR24-2(T) | 98 | 57 |
| Pi19 | P02-B3 | 10 days | random | No | 859 | Burkholderia sordidicola S5-B(T) | 99 | 59 |
| Pi19 | P02-B4 | 10 days | random | No | 904 | Burkholderia udeis LMG 27134(T) | 99 | 62 |
| Pi19 | P02-B5 | 10 days | random | No | 585 | Burkholderia sordidicola S5-B(T) | 98 | 40 |
| Pi19 | P02-C1 | 10 days | random | No | 868 | Mycobacterium austroafricanum ATCC 33464(T) | 98 | 60 |
| Pi19 | P02-C2 | 10 days | random | No | 906 | Mycobacterium vanbaalenii PYR-1(T) | 98 | 63 |
| Pi19 | P02-C3 | 10 days | random | No | 835 | Burkholderia udeis LMG 27134(T) | 99 | 58 |
| Pi19 | P02-C4 | 10 days | random | No | 861 | Burkholderia udeis LMG 27134(T) | 99 | 59 |
| Pi19 | P02-C5 | 10 days | random | No | 781 | Burkholderia phenazinium LMG 2247(T) | 99 | 54 |
| Pi19 | P02-C6 | 10 days | random | No | 852 | Gryllotalpicola daejeonensis RU-04(T) | 99 | 59 |
| Pi19 | P02-D1 | 10 days | random | No | 871 | Sphingomonas ginsenosidimutans Gsoil 1429(T) | 97 | 62 |
| Pi19 | P02-D2 | 10 days | random | No | 838 | Rhodanobacter ginsengisoli GR17-7(T) | 99 | 57 |
| Pi19 | P02-D3 | 10 days | random | No | 766 | Gryllotalpicola daejeonensis RU-04(T) | 99 | 53 |
| Pi19 | P02-D4 | 10 days | random | No | 602 | Burkholderia sordidicola S5-B(T) | 99 | 41 |
| Pi19 | P02-D5 | 10 days | random | No | 782 | Burkholderia sordidicola S5-B(T) | 99 | 54 |
| Pi19 | P02-D6 | 10 days | random | No | 367 | Rhodanobacter ginsenosidimutans Gsoil 3054(T) | 99 | 25 |
| Pi19 | P24-A1 | 3 weeks | random | No | 920 | Gryllotalpicola daejeonensis RU-04(T) | 99 | 64 |
| Pi19 | P24-A2 | 3 weeks | random | No | 1000 | Rhodanobacter umsongensis GR24-2(T) | 99 | 68 |
| Pi19 | P24-A3 | 3 weeks | random | No | 832 | Mycobacterium austroafricanum ATCC 33464(T) | 99 | 58 |
| Pi19 | P24-A4 | 3 weeks | random | No | 654 | Mycobacterium austroafricanum ATCC 33464(T) | 97 | 45 |
| Pi19 | P24-A5 | 3 weeks | random | No | 482 | Mycobacterium austroafricanum ATCC 33464(T) | 98 | 33 |
| Pi19 | P24-A6 | 3 weeks | random | No | 525 | Rhodanobacter umsongensis GR24-2(T) | 99 | 36 |
| Pi19 | P24-B1 | 3 weeks | random | No | 695 | Rhodanobacter umsongensis GR24-2(T) | 99 | 47 |
| Pi19 | P24-B2 | 3 weeks | random | No | 812 | Labedella gwakjiensis KSW2-17(T) | 100 | 56 |
| Pi19 | P24-B4 | 3 weeks | random | No | 897 | Streptacidiphilus durhamensis FSCA67(T) | 99 | 62 |
| Pi19 | P24-B5 | 3 weeks | random | No | 914 | Methylovirgula ligni BW863(T) | 99 | 65 |
| Pi19 | P24-B6 | 3 weeks | random | No | 978 | Acidisphaera rubrifaciens HS-AP3(T) | 95 | 69 |
| Pi19 | P24-C1 | 3 weeks | random | No | 1022 | Granulicella pectinivorans TPB6011(T) | 97 | 72 |

| | | | | | | | | |
|---|---|---|---|---|---|---|---|---|
| Pi19 | P24-C2 | 3 weeks | random | No | 665 | Rhodanobacter umsongensis GR24-2(T) | 99 | 45 |
| Pi19 | P24-C4 | 3 weeks | random | No | 938 | Methylocella tundrae T4(T) | 97 | 67 |
| Pi19 | P24-C6 | 3 weeks | random | No | 771 | Beijerinckia derxii supsp. derxii DSM 2328(T) | 98 | 55 |
| Pi19 | P24-D1 | 3 weeks | random | No | 1073 | Acidicapsa borealis KA1(T) | 98 | 76 |
| Pi19 | P24-D2 | 3 weeks | random | No | 997 | Dyella jiangningensis SBZ3-12(T) | 99 | 68 |
| Pi19 | P24-D3 | 3 weeks | random | No | 1061 | 'Acidipila rosea' AP8(T) | 98 | 75 |
| Pi19 | P24-D4 | 3 weeks | random | No | 631 | Dyella kyungheensis THG-B117(T) | 99 | 43 |
| Pi19 | P24-D5 | 3 weeks | random | No | 1066 | Labedella gwakjiensis KSW2-17(T) | 99 | 74 |
| Pi19 | P24-D6 | 3 weeks | random | No | 1030 | Acidisphaera rubrifaciens HS-AP3(T) | 96 | 73 |
| Pi19 | P43-A1 | 6 weeks | random | No | 1014 | Mycobacterium abscessus subsp. bolletii BD(T) | 100 | 70 |
| Pi19 | P43-A2 | 6 weeks | random | No | 974 | Jatrophihabitans endophyticus S9650(T) | 98 | 68 |
| Pi19 | P43-A3 | 6 weeks | random | No | 1101 | Mycobacterium rhodesiae DSM 44223(T) | 99 | 76 |
| Pi19 | P43-A5 | 6 weeks | random | No | 896 | Acidisoma tundrae WM1(T) | 98 | 63 |
| Pi19 | P43-A6 | 6 weeks | random | No | 939 | Jatrophihabitans endophyticus S9650(T) | 100 | 65 |
| Pi19 | P43-B1 | 6 weeks | random | No | 968 | Edaphobacter modestus Jbg-1(T) | 97 | 68 |
| Pi19 | P43-B2 | 6 weeks | random | No | 932 | Edaphobacter modestus Jbg-1(T) | 97 | 65 |
| Pi19 | P43-B3 | 6 weeks | random | No | 1015 | Mycobacterium austroafricanum ATCC 33464(T) | 99 | 70 |
| Pi19 | P43-B4 | 6 weeks | random | No | 850 | Acidicapsa borealis KA1(T) | 98 | 60 |
| Pi19 | P43-B5 | 6 weeks | random | No | 999 | Acidisphaera rubrifaciens HS-AP3(T) | 96 | 71 |
| Pi20 | P09-A1 | 10 days | morphology | No | 495 | Burkholderia phenazinium LMG 2247(T) | 99 | 34 |
| Pi20 | P09-A2 | 10 days | morphology | No | 1024 | Dyella japonica XD53(T) | 100 | 70 |
| Pi20 | P09-A5 | 10 days | random | No | 1068 | Rhodanobacter terrae GP18-1(T) | 99 | 73 |
| Pi20 | P09-A6 | 10 days | random | No | 893 | Rhodanobacter ginsengisoli GR17-7(T) | 99 | 61 |
| Pi20 | P09-B1 | 10 days | random | No | 657 | Burkholderia phenazinium LMG 2247(T) | 99 | 45 |
| Pi20 | P09-B2 | 10 days | random | No | 974 | Mycobacterium phocaicum CIP 108542(T) | 99 | 68 |
| Pi20 | P09-B4 | 10 days | random | No | 1113 | Rhodanobacter umsongensis GR24-2(T) | 99 | 76 |
| Pi20 | P09-B5 | 10 days | random | No | 116 | Rhodanobacter panaciterrae LnR5-47(T) | 100 | 8 |
| Pi20 | P09-B6 | 10 days | random | No | 1026 | Burkholderia sordidicola S5-B(T) | 98 | 71 |
| Pi20 | P09-C1 | 10 days | random | No | 1170 | Rudaeicoccus suwonensis HOR6-4(T) | 96 | 81 |
| Pi20 | P09-C2 | 10 days | random | No | 1049 | Rudaeicoccus suwonensis HOR6-4(T) | 96 | 72 |
| Pi20 | P09-C3 | 10 days | random | No | 821 | Burkholderia graminis C4D1M(T) | 99 | 57 |
| Pi20 | P09-C4 | 10 days | random | No | 1018 | Rhodanobacter fulvus Jip2(T) | 99 | 69 |
| Pi20 | P09-C5 | 10 days | random | No | 970 | Streptacidiphilus durhamensis FSCA67(T) | 100 | 67 |

| | | | | | | | | |
|---|---|---|---|---|---|---|---|---|
| Pi20 | P09-C6 | 10 days | random | No | 762 | Agreia pratensis P 229/10(T) | 99 | 53 |
| Pi20 | P09-D1 | 10 days | random | No | 1049 | Dyella ginsengisoli Gsoil 3046(T) | 99 | 72 |
| Pi20 | P09-D2 | 10 days | random | No | 951 | Frondihabitans sucicola GRS42(T) | 100 | 66 |
| Pi20 | P09-D3 | 10 days | random | No | 652 | Humibacter antri D7-27(T) | 97 | 45 |
| Pi20 | P09-D4 | 10 days | random | No | 924 | Luteimicrobium album RI148-Li105(T) | 100 | 64 |
| Pi20 | P09-D6 | 10 days | random | No | 578 | Burkholderia phenazinium LMG 2247(T) | 98 | 40 |
| Pi20 | P29-A1 | 3 weeks | random | No | 992 | Rhodanobacter ginsengisoli GR17-7(T) | 98 | 68 |
| Pi20 | P29-A2 | 3 weeks | random | No | 918 | Mycobacterium fluoranthenivorans FA-4(T) | 99 | 64 |
| Pi20 | P29-A3 | 3 weeks | random | No | 919 | Mycobacterium austroafricanum ATCC 33464(T) | 99 | 64 |
| Pi20 | P29-A4 | 3 weeks | random | No | 520 | Methylovirgula ligni BW863(T) | 96 | 37 |
| Pi20 | P29-A5 | 3 weeks | random | No | 1008 | Mycobacterium neoaurum ATCC 25795(T) | 98 | 69 |
| Pi20 | P29-A6 | 3 weeks | random | No | 923 | Mycobacterium neoaurum ATCC 25795(T) | 99 | 64 |
| Pi20 | P29-B1 | 3 weeks | random | No | 422 | Rhodanobacter ginsengisoli GR17-7(T) | 100 | 29 |
| Pi20 | P29-B2 | 3 weeks | random | No | 422 | Rhodanobacter ginsengisoli GR17-7(T) | 99 | 29 |
| Pi20 | P29-B4 | 3 weeks | random | No | 859 | Marmoricola aequoreus SST-45(T) | 99 | 60 |
| Pi20 | P29-B5 | 3 weeks | random | No | 703 | Actinospica robiniae GE134769(T) | 99 | 49 |
| Pi20 | P29-B6 | 3 weeks | random | No | 984 | Methylosinus trichosporium OB3b(T) | 95 | 70 |
| Pi20 | P29-C1 | 3 weeks | random | No | 647 | Mycobacterium austroafricanum ATCC 33464(T) | 99 | 45 |
| Pi20 | P29-C2 | 3 weeks | random | No | 999 | Rhodanobacter ginsengisoli GR17-7(T) | 99 | 68 |
| Pi20 | P29-C4 | 3 weeks | random | No | 899 | Roseiarcus fermentans Pf56(T) | 99 | 64 |
| Pi20 | P29-C5 | 3 weeks | random | No | 982 | Rhodanobacter ginsengisoli GR17-7(T) | 99 | 67 |
| Pi20 | P29-C6 | 3 weeks | random | No | 355 | Actinospica acidiphila GE134766(T) | 100 | 24 |
| Pi20 | P29-D1 | 3 weeks | random | No | 977 | Rhodanobacter ginsengisoli GR17-7(T) | 99 | 67 |
| Pi20 | P29-D2 | 3 weeks | random | No | 895 | Mycobacterium neoaurum ATCC 25795(T) | 99 | 62 |
| Pi20 | P29-D3 | 3 weeks | random | No | 781 | Roseiarcus fermentans Pf56(T) | 99 | 56 |
| Pi20 | P29-D4 | 3 weeks | random | No | 736 | Humibacter antri D7-27(T) | 98 | 51 |
| Pi20 | P29-D5 | 3 weeks | random | No | 898 | Rhodococcus wratislaviensis NBRC 100605(T) | 97 | 63 |
| Pi20 | P45-A1 | 6 weeks | random | No | 1006 | Mycobacterium abscessus subsp, bolletii BD(T) | 100 | 70 |
| Pi20 | P45-A2 | 6 weeks | random | No | 629 | Methylovirgula ligni BW863(T) | 98 | 45 |
| Pi20 | P45-A3 | 6 weeks | random | No | 988 | Methylovirgula ligni BW863(T) | 99 | 70 |
| Pi20 | P45-A4 | 6 weeks | random | No | 342 | Mycobacterium abscessus subsp, bolletii BD(T) | 99 | 24 |
| Pi20 | P45-A5 | 6 weeks | random | No | 780 | Ancylobacter dichloromethanicus DM16(T) | 94 | 56 |
| Pi20 | P45-B1 | 6 weeks | random | No | 557 | Pseudonocardia acaciae DSM 45401(T) | 98 | 39 |

Suppl. Table 6: Bacterial community composition in decaying pine wood, based on the relative abundance of randomly picked isolates.

| Phylum | Family | Genus | Pi04 | Pi05 | Pi06 | Pi08 | Pi09 | Pi11 | Pi12 | Pi13 | Pi14 | Pi16 | Pi17 |
|---|---|---|---|---|---|---|---|---|---|---|---|---|---|
| Acidobacteria (Gp1) | | Acidicapsa | 0% | 0% | 0% | 0% | 0% | 0% | 0% | 0% | 0% | 0% | 0% |
| | | Acidipila | 0% | 0% | 0% | 0% | 0% | 9% | 1% | 0% | 0% | 1% | 0% |
| | | Granulicella | 0% | 0% | 0% | 0% | 0% | 7% | 5% | 2% | 0% | 3% | 5% |
| | | Terriglobus | 0% | 5% | 0% | 0% | 0% | 0% | 0% | 0% | 0% | 0% | 0% |
| | | unclassified | 0% | 0% | 0% | 0% | 0% | 5% | 0% | 0% | 0% | 0% | 0% |
| Actinobacteria | Actinospicaceae | Actinospica | 0% | 0% | 0% | 0% | 0% | 0% | 0% | 0% | 0% | 0% | 0% |
| | Cellulomonadaceae | Cellulomonas | 0% | 0% | 0% | 0% | 0% | 0% | 0% | 0% | 1% | 0% | 0% |
| | Microbacteriaceae | Frondihabitans | 0% | 0% | 0% | 0% | 85% | 0% | 0% | 0% | 0% | 0% | 0% |
| | Microbacteriaceae | Leifsonia | 0% | 0% | 0% | 0% | 0% | 0% | 0% | 0% | 0% | 0% | 0% |
| | Microbacteriaceae | Gryllotalpicola sp. | 0% | 0% | 13% | 11% | 0% | 0% | 0% | 0% | 3% | 0% | 0% |
| | Microbacteriaceae | Curtobacterium | 0% | 0% | 0% | 0% | 0% | 0% | 0% | 0% | 0% | 1% | 0% |
| | Microbacteriaceae | Microbacterium | 0% | 0% | 0% | 0% | 0% | 0% | 0% | 1% | 0% | 0% | 0% |
| | Microbacteriaceae | unclassified | 0% | 0% | 15% | 0% | 6% | 0% | 3% | 8% | 8% | 1% | 15% |
| | Mycobacteriaceae | Mycobacterium | 0% | 0% | 0% | 0% | 0% | 0% | 0% | 19% | 0% | 0% | 9% |
| | Nocardiaceae | Rhodococcus | 0% | 0% | 0% | 0% | 0% | 0% | 0% | 0% | 0% | 0% | 0% |
| | Nocardioidaceae | Marmoricola | 0% | 0% | 0% | 0% | 0% | 0% | 0% | 2% | 0% | 0% | 0% |
| | Pseudonocardiaceae | Pseudonocardia | 0% | 0% | 0% | 0% | 0% | 0% | 0% | 0% | 0% | 0% | 3% |
| | Streptomycetaceae | Kitasatospora | 0% | 0% | 0% | 0% | 0% | 0% | 0% | 0% | 0% | 0% | 3% |
| | Streptomycetaceae | Streptacidiphilus | 0% | 0% | 0% | 0% | 0% | 0% | 0% | 0% | 1% | 0% | 7% |
| | unclassified Actinomycetales | unclassified | 0% | 0% | 0% | 0% | 0% | 0% | 0% | 0% | 0% | 0% | 7% |
| Proteobacteria (Alpha) | Acetobacteraceae | Acidisoma | 0% | 0% | 0% | 0% | 0% | 3% | 4% | 3% | 10% | 6% | 0% |
| | Acetobacteraceae | Acidisphaera | 0% | 0% | 0% | 0% | 0% | 0% | 3% | 0% | 0% | 1% | 0% |
| | Acetobacteraceae | unclassified | 0% | 5% | 15% | 0% | 0% | 15% | 42% | 4% | 20% | 4% | 14% |
| | Beijerinckiaceae | Beijerinckia | 0% | 0% | 0% | 6% | 0% | 5% | 0% | 0% | 0% | 0% | 0% |
| | Beijerinckiaceae | Methylocella | 0% | 0% | 0% | 0% | 0% | 5% | 0% | 0% | 0% | 0% | 0% |
| | Beijerinckiaceae | Methylovirgula | 0% | 0% | 0% | 39% | 0% | 18% | 21% | 22% | 20% | 0% | 20% |
| | Beijerinckiaceae | unclassified | 0% | 0% | 0% | 3% | 0% | 16% | 3% | 8% | 0% | 0% | 0% |
| | Bradyrhizobiaceae | Bradyrhizobium | 0% | 0% | 0% | 0% | 0% | 3% | 0% | 6% | 13% | 0% | 0% |
| | Caulobacteraceae | Phenylobacterium | 0% | 0% | 13% | 0% | 0% | 10% | 0% | 4% | 0% | 0% | 0% |
| | Caulobacteraceae | unclassified | 0% | 0% | 0% | 0% | 0% | 5% | 0% | 0% | 0% | 0% | 0% |

| Phylum | Family | Genus | Pi18 | Pi19 | Pi20 | Average |
|---|---|---|---|---|---|---|
| Acidobacteria (Gp1) | | Acidicapsa | 0% | 7% | 0% | 0% |
| | | Acidipila | 0% | 2% | 0% | 1% |
| | | Granulicella | 0% | 12% | 0% | 2% |
| | | Terriglobus | 0% | 0% | 0% | 0% |
| | | unclassified | 0% | 0% | 0% | 0% |
| Actinobacteria | Actinospicaceae | Actinospica | 1% | 0% | 3% | 0% |
| | Cellulomonadaceae | Cellulomonas | 0% | 0% | 2% | 0% |
| | Microbacteriaceae | Frondihabitans | 0% | 0% | 2% | 6% |
| | Microbacteriaceae | Leifsonia | 0% | 0% | 2% | 0% |
| | Microbacteriaceae | Gryllotalpicola sp. | 0% | 4% | 0% | 2% |
| | Microbacteriaceae | Curtobacterium | 0% | 0% | 0% | 0% |
| | Microbacteriaceae | Microbacterium | 0% | 0% | 0% | 0% |
| | Microbacteriaceae | unclassified | 0% | 3% | 4% | 5% |
| | Mycobacteriaceae | Mycobacterium | 36% | 22% | 21% | 8% |
| | Nocardiaceae | Rhodococcus | 0% | 0% | 1% | 0% |
| | Nocardioidaceae | Marmoricola | 0% | 0% | 1% | 0% |
| | Pseudonocardiaceae | Pseudonocardia | 0% | 0% | 3% | 0% |
| | Streptomycetaceae | Kitasatospora | 0% | 0% | 0% | 0% |
| | Streptomycetaceae | Streptacidiphilus | 6% | 2% | 2% | 1% |
| | unclassified Actinomycetales | unclassified | 0% | 10% | 8% | 2% |
| Proteobacteria (Alpha) | Acetobacteraceae | Acidisoma | 0% | 5% | 0% | 2% |
| | Acetobacteraceae | Acidisphaera | 0% | 0% | 0% | 0% |
| | Acetobacteraceae | unclassified | 19% | 8% | 0% | 10% |
| | Beijerinckiaceae | Beijerinckia | 0% | 2% | 0% | 1% |
| | Beijerinckiaceae | Methylocella | 0% | 2% | 0% | 0% |
| | Beijerinckiaceae | Methylovirgula | 0% | 2% | 7% | 11% |
| | Beijerinckiaceae | unclassified | 19% | 0% | 3% | 4% |
| | Bradyrhizobiaceae | Bradyrhizobium | 0% | 0% | 0% | 2% |
| | Caulobacteraceae | Phenylobacterium | 0% | 0% | 0% | 2% |
| | Caulobacteraceae | unclassified | 0% | 0% | 0% | 0% |
